# MetaSplice: pathogenicity prediction and ACMG/AMP evidence for intronic variants outside the canonical splice sites

**DOI:** 10.64898/2026.07.23.740393

**Authors:** Xiaoming Liu

## Abstract

Intronic variants outside the canonical splice-site dinucleotides are difficult to interpret: splicing predictors are least accurate there, and clinical classification requires evidence about pathogenicity rather than predicted splicing alone. We developed MetaSplice, a gradient-boosted ensemble that combines SpliceTransformer and Pangolin splicing predictions with conservation, exon context and gene-level constraint, without allele frequency, to rank intronic single-nucleotide variants by pathogenicity. We trained it on 413,199 variants from the August 2025 ClinVar release and tested it on 46,112 variants classified as (likely) pathogenic or (likely) benign in September 2026 but not in August 2025. Among 40,296 non-canonical variants scored by both tools, MetaSplice reached an area under the precision–recall curve of 0.919, versus 0.757 for SpliceAI. On all 40,619 non-canonical variants, it reached 0.916, versus 0.364 for CADD. The advantage was largest at deep intronic positions, persisted when each variant was scored by a model trained without its gene, and was not clear in the polypyrimidine tract. Against experimentally assayed splicing, MetaSplice detected 92.8% of splice-altering variants at a 1.2% false-positive rate. From likelihood ratios calibrated against ClinVar, we derived a candidate rule that assigns PP3 to scores of 0.544 or higher and BP4 to scores of 0.0017 or lower, both at Supporting strength. In the test set, this rule gave PP3 to 0.20% of 37,263 benign variants, versus 4.44% under the ClinGen-recommended SpliceAI thresholds, and to 1.0% versus 31.2% of 706 formerly uncertain variants since classified benign. MetaSplice complements splicing predictors, and its candidate PP3/BP4 rule warrants independent evaluation.

## Introduction

Variants that disrupt pre-mRNA splicing account for an estimated 15% to one third of disease-causing mutations.^1,2^ Single-nucleotide variants (SNVs) at the canonical GT–AG dinucleotides at intron ends are usually interpreted as loss-of-function variants.^3^ Variants elsewhere in the intron, in the extended donor and acceptor regions, the polypyrimidine tract (PPT), the branch-point region and deeper in the intron, are harder to interpret, because their effects on splicing are less predictable, and many remain variants of uncertain significance (VUS) in ClinVar.^4^

Two kinds of computational tool address these variants. Splicing predictors estimate how a variant changes splicing. Deep-learning models such as SpliceAI, Pangolin and SpliceTransformer predict changes in splice-site usage from sequence and are widely used in variant interpretation.^5–7^ SPIDEX predicts changes in exon inclusion.^8^ Their accuracy is highest at the canonical sites and declines with distance from the exon boundary, and a predicted splicing change does not by itself establish pathogenicity, which also depends on the gene, the transcript and the consequence of the altered RNA. Pathogenicity predictors instead learn from classified variants. Our earlier ensemble, dbscSNV, combined several splice-site scores to identify splice-altering variants but covers only the splicing consensus region (−3 to +8 at the 5′ splice site and −12 to +2 at the 3^′^ splice site).^9^ S-CAP trained region-specific models to predict the pathogenicity of rare splice-region variants,^10^ SQUIRLS uses interpretable sequence features to prioritize splice variants in diagnostic sequencing,^11^ and CADD integrates many annotations, including SpliceAI and MMSplice scores, into a genome-wide deleteriousness score.^12,13^ Conservation scores capture evolutionary constraint but lack splice-specific resolution.^14,15^ It remains unclear how much gene and conservation context adds to current splicing predictions when the target is clinical pathogenicity at every intronic position.

Clinical use of such scores also requires calibrated evidence strengths. Under the guidelines of the American College of Medical Genetics and Genomics and the Association for Molecular Pathology (ACMG/AMP), computational predictions provide supporting evidence for (PP3) or against (BP4) pathogenicity.^3^ Expressing each evidence strength as a likelihood ratio^16^ has allowed the Clinical Genome Resource (ClinGen) to calibrate missense predictors for PP3 up to Strong and for BP4 up to Moderate strength.^17,18^ For splicing, the ClinGen Sequence Variant Interpretation Splicing Subgroup recommends PP3 for SpliceAI scores of 0.2 or higher and BP4 for scores of 0.1 or lower, applied at Supporting strength.^19^ ClinGen does not calibrate predictors that use allele frequency, because frequency is assessed by separate criteria.^18^

Here we present MetaSplice, a gradient-boosted ensemble of 38 features that combines SpliceTransformer and Pangolin predictions with conservation, exon context and gene-level constraint to score intronic SNVs by pathogenicity, without using allele frequency. We trained it on ClinVar and compared it with SpliceAI and CADD on variants classified after the training release, with emphasis on variants outside the canonical sites. We tested whether its advantage depends on the variant’s gene, evaluated it against experimentally assayed and literature-confirmed splice-altering variants, and calibrated its scores for PP3 and BP4, comparing a candidate rule with the ClinGen SpliceAI rule.

## Material and methods

### Variant sets

We used the GRCh38 ClinVar^4^ releases of 31 August 2025 for training and 28 September 2026 for testing. SNVs classified pathogenic or likely pathogenic (P/LP) were labeled positive and those classified benign or likely benign (B/LB) negative. VUS and conflicting classifications were not used as labels. The scope was the intronic positions of GENCODE v50 transcripts tagged MANE Select or MANE Plus Clinical (19,373 transcripts in 19,299 genes).^20,21^ Each variant was assigned to one of seven regions by its distance from the nearest exon boundary of its gene (Table 1): the canonical donor and acceptor sites (intronic positions +1 to +2 and −1 to−2), the donor region (+3 to +8), the acceptor region (−3 to −10), the PPT region (−11 to −25), the branch-point region (−26 to −44) and deep intronic positions beyond these windows. The PPT and branch-point regions are positional windows, not mapped motifs. A position near more than one boundary took the first applicable region in the order canonical acceptor, canonical donor, acceptor, donor, PPT, branch point and deep intronic. Deep intronic positions inside an exon of any GENCODE v50 basic transcript were out of scope, and a deep-intronic sensitivity set further excluded positions that are canonical or exonic in any GENCODE v50 transcript. We call the five regions other than the canonical sites non-canonical.

**Table 1.** ClinVar variant sets by region. Number of variants in each region, with P/LP variants in parentheses.

| Region | Intronic position<sup>a</sup> | Training (ClinVar 2025-08-31) | Temporal test set (ClinVar 2026-09-28)<sup>b</sup> |
| --- | --- | --- | --- |
| Canonical donor site | +1 to +2 | 20,576 (20,443) | 2,951 (2,932) |
| Canonical acceptor site | −1 to −2 | 17,126 (16,990) | 2,542 (2,522) |
| Donor region | +3 to +8 | 26,476 (1,862) | 3,314 (283) |
| Acceptor region | −3 to −10 | 64,628 (784) | 7,021 (123) |
| PPT region | −11 to −25 | 86,776 (348) | 12,410 (61) |
| Branch-point region | −26 to −44 | 5,822 (53) | 395 (9) |
| Deep intronic | beyond +8 and −44 | 191,795 (552) | 17,479 (126) |
| Deep-intronic sensitivity set<sup>c</sup> |  | 159,226 (364) | 14,459 (99) |
| All regions |  | 413,199 (41,032) | 46,112 (6,056) |
| Non-canonical |  | 375,497 (3,599) | 40,619 (602) |
| Calibration cohort, ≥ 1 review star<sup>d</sup> |  | 294,612 (3,018) | 37,841 (578) |

<sup>a</sup>Distance into the intron from the nearest exon boundary of a GENCODE v50 MANE Select or MANE Plus Clinical transcript: + downstream of a donor, − upstream of an acceptor. A position near more than one boundary takes the first region that applies in the order canonical acceptor, canonical donor, acceptor, donor, PPT, branch point, deep intronic. <sup>b</sup>In-scope SNVs classified P/LP or B/LB in ClinVar on 28 September 2026 that were not classified P/LP or B/LB in the training release. SpliceAI scores 45,749 test variants (6,015 P/LP), including 40,296 non-canonical (600 P/LP), and CADD scores all of them. <sup>c</sup>Deep intronic positions that are neither canonical nor exonic in any GENCODE v50 transcript. <sup>d</sup>Non-canonical variants with gnomAD allele frequency below 0.01 in genes with at least one P/LP variant in the training release.
P/LP, pathogenic or likely pathogenic; B/LB, benign or likely benign; PPT, polypyrimidine tract; SNV, single-nucleotide variant.

The training set comprised 413,199 SNVs (41,032 P/LP) in 6,961 genes (Table 1). The temporal test set comprised in-scope SNVs classified P/LP or B/LB in the September 2026 release that had not been classified P/LP or B/LB in the training release, matched by allele or ClinVar identifier. It contained 46,112 variants (6,056 P/LP), including 40,619 non-canonical variants (602 P/LP).

Four further sets addressed specific questions (Table S1). First, resolved VUS were variants that were uncertain or conflicting in the training release and classified P/LP or B/LB in September 2026. Second, remaining VUS were uncertain or conflicting variants not classified P/LP or B/LB by September 2026 (Supplemental Methods). Third, to test recovery of splice-altering variants identified independently of ClinVar, we assembled intronic variants reported as splice-altering with experimental support (RT-PCR, minigene or other functional assays). Candidates were extracted from PubMed abstracts with a large language model (Claude Opus 4.6, Anthropic), and a blinded manual review of 50 randomly selected candidates found 49 fully and one partially supported. After mapping to GRCh38, scope filtering and removal of 30 training variants, 697 variants remained, all of them positives. Fourth, to evaluate MetaSplice against experimentally assayed splicing, we used SpliceVarDB,^22^ which classifies variants by experimental splicing assays as splice-altering, low-frequency splice-altering or normal. We rebuilt these classifications by applying SpliceVarDB’s published criteria to the data of its three largest source studies, MFASS,^23^ SAVNet^24^ and MiSplice,^25^ and of four literature studies (Supplemental Methods). After mapping, scope filtering and removal of training variants, 22,314 unique SNVs remained, from five of these studies. After excluding 40 whose assays disagreed, 22,274 remained. Mapping and filtering are described in the Supplemental Methods.

### Features and model

MetaSplice uses 38 features (Table S2). The splicing features are the aggregate and 15 tissue-specific SpliceTransformer scores^7^ and the largest predicted increase, decrease and absolute change in splice-site usage from Pangolin.^6^ Conservation is represented by phyloP and phastCons scores from the 100-way vertebrate and 470-way mammalian alignments.^14,15^ The other features are the DANN^26^ and GPN-MSA^27^ variant scores, three gene-level annotations from dbNSFP 5.4^28^ (the gnomAD probability of loss-of-function intolerance [pLI], the loss-of-function observed/expected upper bound fraction [LOEUF] and the ClinGen haploinsufficiency score; rebuilt from gnomAD, ClinGen and HGNC with the same rules), three exon-context features (the length of the nearest exon, that length modulo three and the distance to its boundary) and seven region indicators. SpliceTransformer and Pangolin were run on GRCh38 with annotations built from the same MANE transcripts. We did not use population allele frequency as an input, so that MetaSplice scores can be combined with the frequency-based ACMG/AMP criteria without counting frequency twice. ClinGen likewise excludes predictors that use allele frequency from calibration.^18^ The same feature pipeline computed the features of every variant set, and its sources and consistency checks are described in the Supplemental Methods.

We trained an XGBoost classifier^29^ with 300 trees, maximum depth 6, learning rate 0.1, histogram-based tree construction and seed 42, with the positive-class weight set to the ratio of B/LB to P/LP variants, missing values handled natively and no hyperparameter tuning. Out-of-fold predictions came from five-fold cross-validation in which all variants of a gene were assigned to the same fold. The final model was trained on all training variants and scored every other variant set. Feature contributions were computed as SHAP values with TreeSHAP^30^ and summarized as the mean absolute value for each feature, summed within feature groups.

### Comparisons and statistical analysis

We compared MetaSplice with SpliceAI,^5^ the splicing predictor whose thresholds ClinGen has calibrated for splicing evidence,^19^ and with CADD v1.7,^12^ a genome-wide pathogenicity score that incorporates splicing predictions.^13^ For SpliceAI, we used the maximum of the four delta scores in its precomputed masked score files, and for CADD, the PHRED-scaled score. dbscSNV was not compared because it scores only the consensus region, and SQUIRLS because its scoring database was unavailable at the time of analysis. Because P/LP variants are a small minority of non-canonical test variants (1.5%), we measured discrimination by the area under the precision–recall curve (auPRC, computed as average precision), which depends on how well the rare positive class is ranked. The area under the receiver operating characteristic curve (AUROC) was a secondary measure. Because canonical-site variants supplied 90.1% of P/LP test variants and every tool ranked them almost perfectly, non-canonical variants were the primary analysis set. Each tool’s auPRC is reported on the variants it scores, and comparisons between two tools used the variants both score.

To estimate the uncertainty of differences in auPRC, we used a paired bootstrap that resampled P/LP and B/LB variants separately, recalculated both tools’ auPRC in each of 1,000 resamples (seed 42) and took percentile 95% confidence intervals (CIs). To allow for correlation among variants of the same gene, we also resampled genes with replacement (1,000 resamples). We describe a difference as clear when its 95% CI excludes zero. Regions with fewer than 20 P/LP variants were reported as counts only. Single-tool auPRC CIs used the same class-stratified bootstrap.

Most test variants were in genes that also contributed training variants. To test whether performance depended on this, we assigned the 7,194 genes of the training and test sets to five folds at random (seed 42), trained one model per fold on training variants from the other four folds and scored the test variants of that fold with it. To remove gene-level information from the comparison, we computed a within-gene concordance index on the non-canonical test variants that all three tools score: across all pairs of a P/LP and a B/LB variant in the same gene, the proportion in which the P/LP variant scores higher, counting ties as one half. Because both variants in each pair come from the same gene, a score with one value per gene, such as a gene-level constraint metric, ties every pair and has a concordance of 0.5. The index therefore measures ranking within genes only. Its CIs used the gene-resampling bootstrap.

To measure what the full model adds over simpler models, we trained two baselines on part of the feature set: XGBoost on the 19 SpliceTransformer and Pangolin features, and logistic regression on the aggregate SpliceTransformer score, the Pangolin maximum and phyloP (100-way). We measured the contribution of the gene-level features and the general variant-effect scores (DANN and GPN-MSA) by retraining the model without each group and without both. The model without both was also retrained with genes held out as above. All models used the training protocol of the full model.

### Calibration and a candidate PP3/BP4 rule

We calibrated MetaSplice and SpliceAI scores with the likelihood-ratio approach of Pejaver et al.^17^ in a calibration cohort of non-canonical variants with gnomAD^31^ allele frequency below 0.01 in genes with at least one P/LP variant in the training release. Allele frequency served only to define this cohort. Thresholds were derived from out-of-fold scores of training variants with at least one ClinVar review star (294,612 variants, 3,018 P/LP), using local score windows and a one-sided bootstrap bound on the likelihood ratio (Supplemental Methods). The likelihood-ratio requirements were 2.41 (Supporting), 5.79 (Moderate), 33.5 (Strong) and 1,124 (Very Strong) for PP3, and their reciprocals for BP4. They correspond to a prior probability of pathogenicity of 0.0441.^16,17^ We used this prior, estimated for missense variants, because an attempt to estimate it for intronic variants from a gnomAD reference sample did not meet our pre-set fit criteria (Supplemental Methods). The thresholds are therefore exploratory. In the test set’s calibration cohort, we estimated the likelihood ratio of each score interval as its share of P/LP variants divided by its share of B/LB variants, with exact 95% CIs combining 97.5% Clopper–Pearson intervals for the two shares.

The candidate rule assigns PP3 at Supporting strength to MetaSplice scores of 0.544 or higher and BP4 at Supporting strength to scores of 0.0017 or lower (rounded values, with exact values in Table S3). These are the derived thresholds for PP3 Strong and BP4 Moderate, applied at Supporting strength as ClinGen recommends for predicted splicing evidence.^19^ As defined, the rule would replace rather than add to a SpliceAI-based PP3 or BP4 for the same variant. When RNA assays show the variant’s effect on splicing, that evidence is captured with the codes ClinGen recommends for observed splicing (PVS1 at an RNA-based strength, or BP7),^19^ and MetaSplice-based PP3 or BP4 would then not be applied.

We compared the rule with the ClinGen SpliceAI rule (PP3 for scores of 0.2 or higher, BP4 for 0.1 or lower),^19^ using SpliceAI 1.3.1 scores computed without masking over ±4,999 nucleotides, the configuration in which those thresholds were calibrated (Supplemental Methods). The comparison used the 37,841 variants with at least one review star in the test set’s calibration cohort (578 P/LP), 859 resolved VUS and 60,776 remaining VUS. We report the share of P/LP and B/LB variants receiving each code, the likelihood ratio of each code with exact 95% CIs, and paired bootstrap differences in shares, with CIs from class-stratified and gene resampling. A variant without a score received no code.

### AI usage

Claude Code (Anthropic; Claude Opus 4.6 and 5.5) assisted with software and analysis-script development, and Claude Opus 4.6 with extraction of literature candidates. The author reviewed, tested and revised the generated code and remains responsible for the analyses and numerical results. The AI tools used in preparing the manuscript text are listed in the generative AI declaration.

## Results

### MetaSplice combines splicing predictions with conservation and gene context

MetaSplice was trained on 413,199 intronic SNVs (41,032 P/LP) in 6,961 genes (Figure 1; Table 1). Canonical-site variants made up 37,433 of the P/LP training variants (91.2%), and P/LP variants were uncommon in the non-canonical regions, from 0.29% of deep intronic variants to 7.0% of donor-region variants. In gene-grouped cross-validation, out-of-fold auPRC was 0.994 for all training variants and 0.878 for the 375,497 non-canonical variants (3,599 P/LP), ranging from 0.956 in the donor region to 0.555 in the branch-point region (Table S4). The Pangolin features contributed most to the model’s predictions, followed by SpliceTransformer, exon context, gene-level annotations and conservation, in both the training and the test variants (Figure 3B).

**Figure 1.**
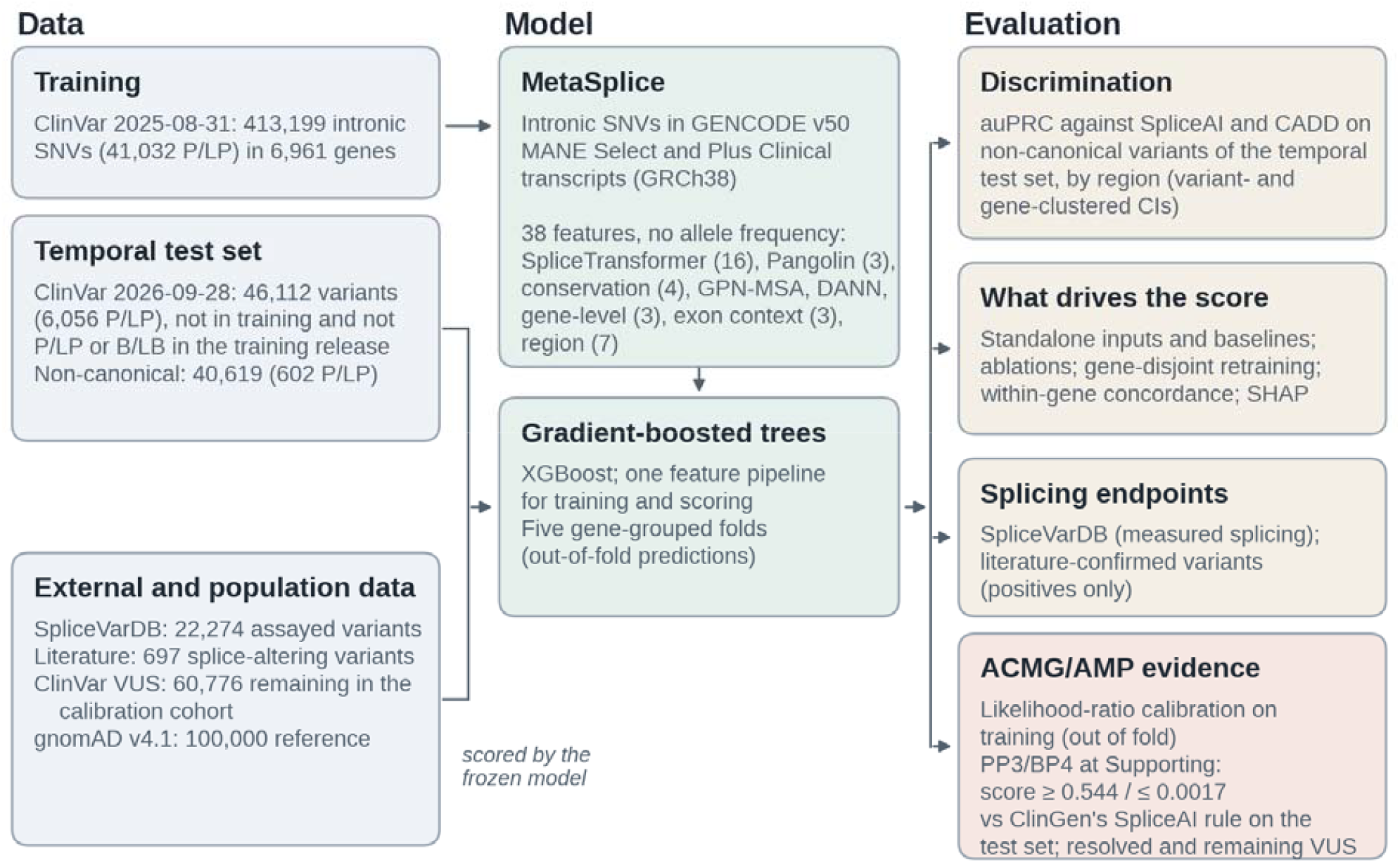
Study design. MetaSplice was trained on intronic single-nucleotide variants classified P/LP or B/LB in ClinVar on 31 August 2025 and evaluated on variants classified P/LP or B/LB in ClinVar on 28 September 2026 but not in the training release, on experimentally assayed (SpliceVarDB) and literature-confirmed splice-altering variants, and for ACMG/AMP evidence. Counts are given in Table 1 and Table S1. P/LP, pathogenic or likely pathogenic; B/LB, benign or likely benign; VUS, variants of uncertain significance.

**Figure 2.**
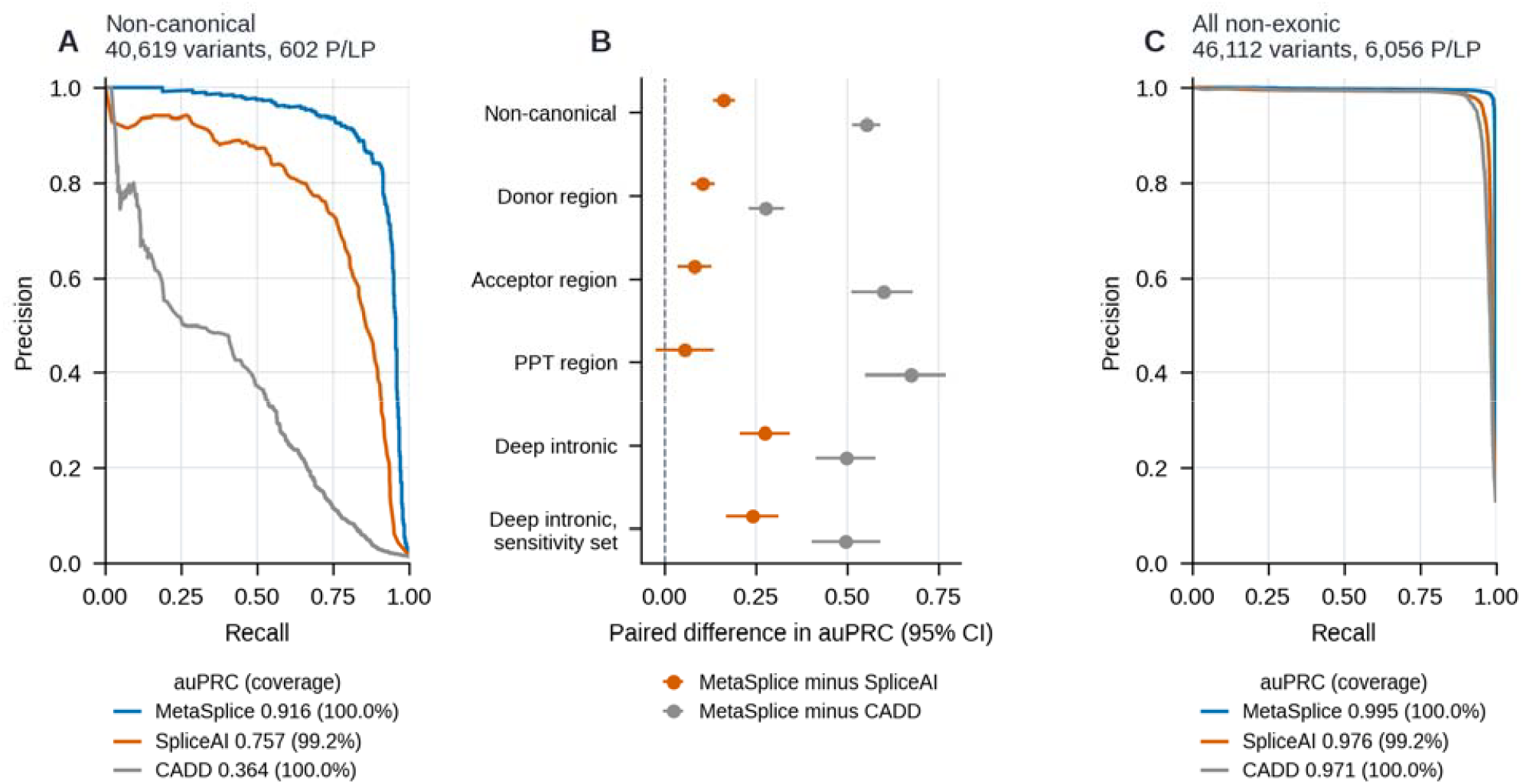
Discrimination on the temporal test set. (A) Precision–recall curves for the 40,619 non-canonical test variants (602 P/LP). Each curve uses the variants the tool scores, and the key gives auPRC and coverage. (B) Paired differences in auPRC between MetaSplice and each comparator by region, on the variants both tools score, with 95% CIs from a class-stratified bootstrap (1,000 resamples). The branch-point region (9 P/LP) is not shown. (C) Precision–recall curves for all 46,112 non-exonic test variants. Canonical-site variants supply 90.1% of P/LP variants. auPRC, area under the precision–recall curve; CI, confidence interval; P/LP, pathogenic or likely pathogenic.

**Figure 3.**
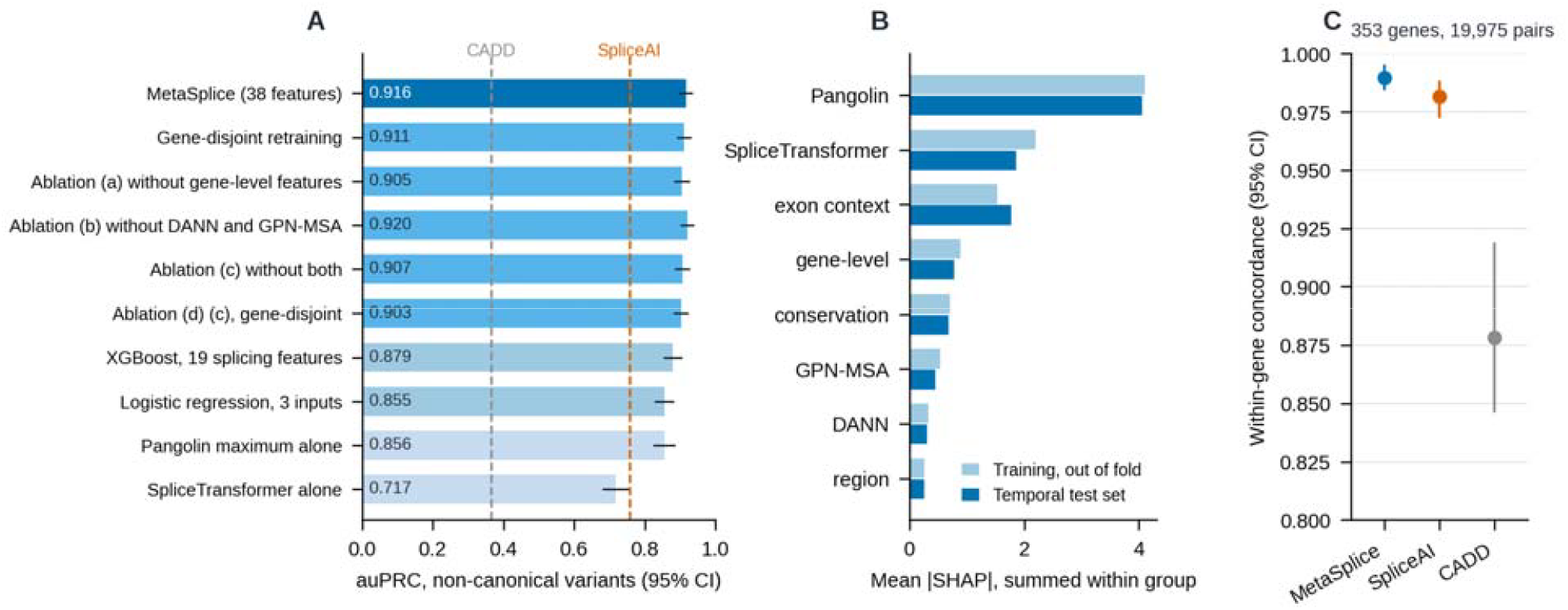
What MetaSplice’s performance depends on. (A) auPRC on the 40,619 non-canonical test variants for MetaSplice, the gene-disjoint model (each variant scored by a model trained without its gene), four ablation models, two baselines and the two strongest single inputs, with 95% bootstrap CIs. Dashed lines mark SpliceAI (on the 99.2% of variants it scores) and CADD. (B) Feature-group importance as mean absolute SHAP value per feature, summed within group, for non-canonical training variants (out-of-fold models) and test variants (full model). (C) Within-gene concordance for the 353 genes with both P/LP and B/LB non-canonical test variants (19,975 pairs) scored by all three tools, with 95% CIs from resampling genes. A score with one value per gene would have a concordance of 0.5. auPRC, area under the precision–recall curve; CI, confidence interval; SHAP, Shapley additive explanations.

### MetaSplice discriminates non-canonical variants better than SpliceAI and CADD

On all 46,112 test variants, auPRC was 0.995 for MetaSplice, 0.976 for SpliceAI and 0.971 for CADD (Figure 2C). These values are dominated by the canonical sites, which supplied 5,454 of the 6,056 P/LP variants (90.1%) but only 39 B/LB variants, and where every tool’s auPRC was at least 0.994 (Table S5). Among the 40,619 non-canonical variants (602 P/LP), auPRC was 0.916 for MetaSplice, 0.757 for SpliceAI and 0.364 for CADD (Figure 2A). On the 40,296 non-canonical variants that SpliceAI scores (600 P/LP), MetaSplice reached 0.919, a paired difference of 0.162 (95% CI 0.134–0.192). Against CADD on all non-canonical variants, the difference was 0.552 (0.512–0.590). With genes resampled, the 95% CIs were 0.128–0.197 (versus SpliceAI) and 0.510–0.595 (versus CADD).

The advantage over SpliceAI differed by region (Figure 2B; Table S5). On the variants both tools score, it was largest at deep intronic positions, where auPRC was 0.824 versus 0.551 (difference 0.273, 95% CI 0.205–0.342), and in the deep-intronic sensitivity set (0.847 versus 0.607; difference 0.240, 95% CI 0.169–0.311). It was smaller in the donor region (0.969 versus 0.865; difference 0.105, 95% CI 0.074–0.138) and the acceptor region (0.947 versus 0.865; difference 0.082, 95% CI 0.036–0.129). In the PPT region the two tools did not clearly differ (0.783 versus 0.728; difference 0.054, 95% CI −0.025 to 0.136). MetaSplice exceeded CADD in every evaluable region. The branch-point region contained only 9 P/LP variants and was not evaluated.

### The ensemble improves on its splicing inputs

To assess how much the ensemble adds to its inputs, we compared MetaSplice with each input alone and with two simpler baselines (Figure 3A; Table S6). Pangolin’s maximum score was the strongest single input (non-canonical auPRC 0.856), followed by the aggregate SpliceTransformer score (0.717), and no other input exceeded 0.281. MetaSplice exceeded Pangolin alone by 0.060 (95% CI 0.039–0.080), an XGBoost model trained on the 19 SpliceTransformer and Pangolin features by 0.037 (0.020–0.053) and a logistic regression on SpliceTransformer, Pangolin and phyloP by 0.061 (0.045–0.076). The gain over Pangolin alone was largest at deep intronic positions (0.120, 95% CI 0.073–0.174) and in the deep-intronic sensitivity set (0.077, 95% CI 0.037–0.123), smaller in the donor region (0.026, 95% CI 0.013– 0.039) and not clear in the acceptor (0.031, 95% CI −0.002 to 0.073) or PPT regions (0.008, 95% CI −0.049 to 0.074).

### The advantage does not depend on having seen the variant’s gene

Of the 40,619 non-canonical test variants, 40,239 (99.1%) were in genes that also contributed training variants. To test whether MetaSplice’s performance depended on this overlap, we scored each test variant with a model trained without its gene. This gene-disjoint model reached a non-canonical auPRC of 0.911 (95% CI 0.893–0.928). The full model’s advantage over it was 0.005 (−0.002 to 0.011), and the gene-disjoint model still exceeded SpliceAI by 0.157 (0.128–0.187) and CADD by 0.548 (0.509–0.586) (Figure 3A). In every evaluable region, the difference between the full and gene-disjoint models had a CI that included zero.

To compare how the tools rank variants of the same gene, we used the within-gene concordance index. In the 353 genes containing both P/LP and B/LB non-canonical test variants (19,975 pairs), concordance was 0.990 (95% CI 0.984–0.995) for MetaSplice, 0.982 (0.972– 0.989) for SpliceAI and 0.878 (0.846–0.919) for CADD (Figure 3C). MetaSplice and SpliceAI did not clearly differ within genes (difference 0.0082, 95% CI −0.0004 to 0.0183), whereas MetaSplice exceeded CADD (difference 0.111, 95% CI 0.075–0.141). MetaSplice and SpliceAI were both close to the maximum of this measure. Because within-gene concordance is a different measure from auPRC and covers only genes with both classes, it does not show how much of the pooled auPRC difference arises within or between genes.

### Removing gene-level or general variant-effect inputs leaves most of the advantage

To test how much MetaSplice depends on its gene-level and general variant-effect inputs, we removed each group separately and then both together (Figure 3A; Table S7). Removing the three gene-level features reduced non-canonical auPRC by 0.011 (95% CI 0.003–0.018). Removing DANN and GPN-MSA caused no clear change (0.004, 95% CI −0.001 to 0.009), and removing both reduced auPRC by 0.009 (0.003–0.017). The model without both groups, retrained with genes held out, reached 0.903 and still exceeded SpliceAI by 0.150 (0.121–0.179). The largest regional reductions were for the gene-level features at deep intronic positions (0.038, 95% CI 0.008–0.074) and for removal of both groups in the PPT region (0.069, 95% CI 0.023– 0.121).

### MetaSplice recovers literature-confirmed and experimentally assayed splice-altering variants

Of the 697 literature-confirmed splice-altering variants, 640 (91.8%) scored 0.1 or higher, including 101 of 130 deep intronic variants (77.7%) (Figure 4C). Because this set has no negatives, it measures sensitivity only.

**Figure 4.**
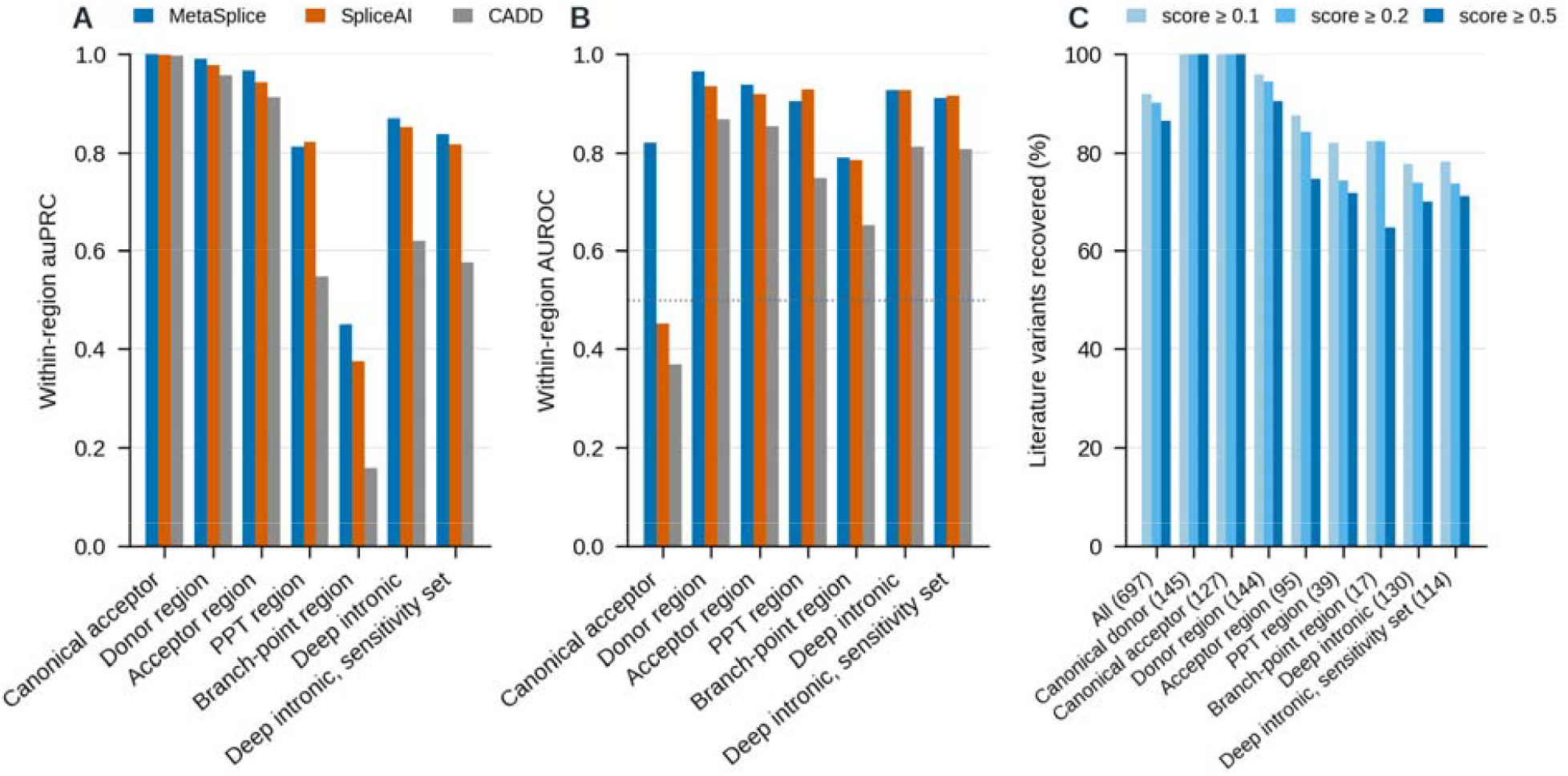
Recovery of experimentally assayed and literature-confirmed splice-altering variants. (A) Within-region auPRC separating SpliceVarDB splice-altering from normal variants (strict endpoint; 12,731 variants), for MetaSplice, SpliceAI and CADD. The canonical donor site, with one normal variant, is omitted. (B) The same comparison by AUROC, which does not depend on the class proportions. The dotted line marks 0.5. (C) Percentage of 697 literature-confirmed splice-altering variants scoring at or above 0.1, 0.2 and 0.5, by region, with numbers of variants in parentheses. auPRC, area under the precision–recall curve; AUROC, area under the receiver operating characteristic curve.

To test whether MetaSplice also identifies variants that alter splicing in experimental assays, we scored the SpliceVarDB set. MetaSplice separated 8,015 experimentally splice-altering variants from 4,716 normal variants with auPRC 0.993, compared with 0.985 for SpliceAI and 0.981 for CADD. A score of 0.1 detected 92.8% of splice-altering variants at a false-positive rate of 1.2%, and a score of 0.5 detected 89.0% at a false-positive rate of 0.7%. Because splice-altering variants are concentrated at canonical sites and normal variants at branch-point and deep intronic positions, within-region comparisons are more informative (Figure 4A–B; Table S8). MetaSplice exceeded SpliceAI in the donor region (auPRC difference 0.013, 95% CI 0.008–0.019) and the acceptor region (0.025, 95% CI 0.015–0.036), whereas in the PPT, branch-point and deep intronic regions the CIs included zero. At deep intronic positions a score of 0.1 detected 66.7% of splice-altering variants at a false-positive rate of 0.3%. When low-frequency splice-altering variants were added to the positives, within-region AUROC fell to 0.48–0.57 for all three tools in the PPT, branch-point and deep intronic regions, where these variants made up 87–96% of the positives.

### A candidate PP3/BP4 rule assigns fewer discordant codes than the ClinGen SpliceAI rule

In the training calibration cohort, out-of-fold MetaSplice scores met the likelihood-ratio requirements for PP3 at 0.052 (Supporting), 0.140 (Moderate), 0.544 (Strong) and 0.994 (Very Strong), and for BP4 at 0.00386 (Supporting), 0.0017 (Moderate) and 0.000275 (Strong) (Table S3). For precomputed SpliceAI scores, the derived PP3 Supporting threshold was 0.2, the value ClinGen recommends,^19^ with Moderate at 0.3 and Strong at 0.49.

To assess the intervals in variants not used to derive them, we estimated each interval’s likelihood ratio in the test set’s calibration cohort. Every MetaSplice interval met its requirement as a point estimate, and its exact 95% CI cleared the requirement for PP3 Very Strong, Strong and Moderate and for BP4 Supporting and Moderate (Figure 5A; Table S9). Two CIs were inconclusive. The PP3 Supporting interval, with 10 P/LP and 179 B/LB variants, had a likelihood ratio of 3.6 (95% CI 1.31–8.38) against a requirement of 2.41, and the BP4 Strong interval had 0.0171 (95% CI 0.00532–0.0407) against a requirement of 0.0298. Every SpliceAI level cleared its requirement.

**Figure 5.**
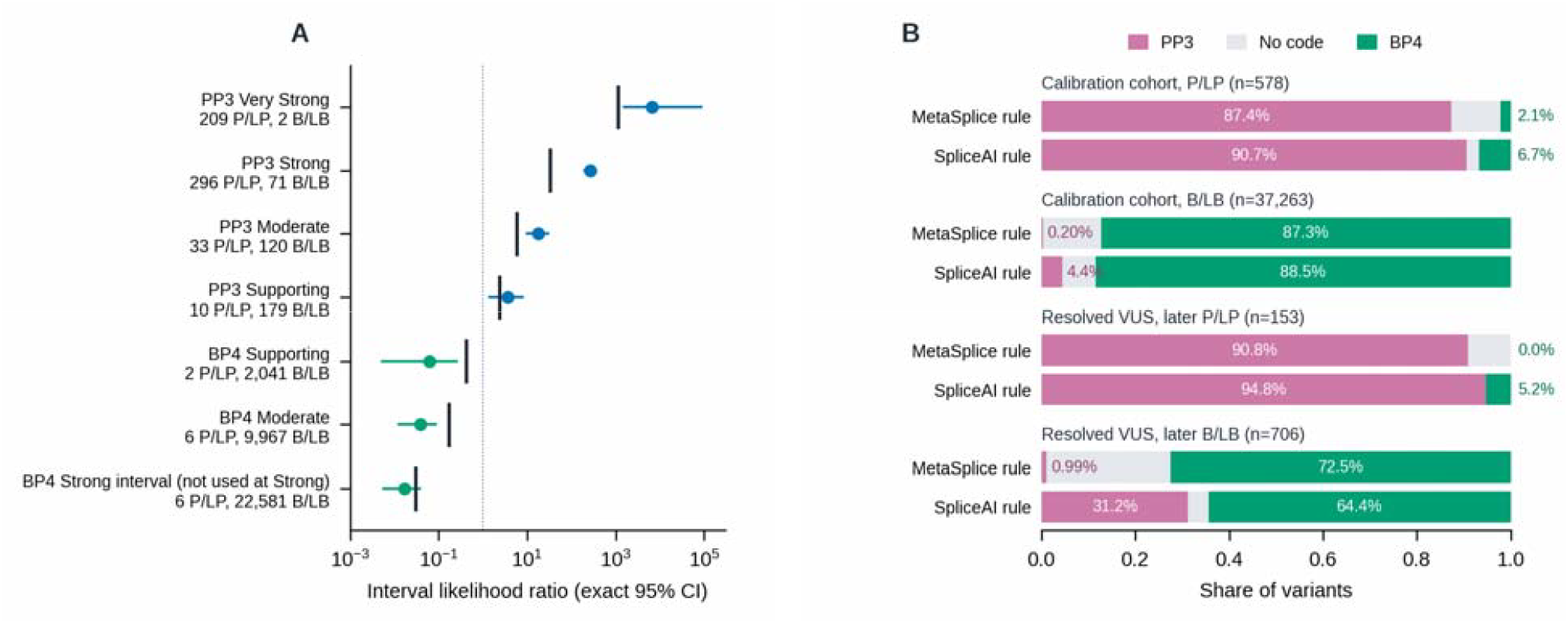
ACMG/AMP evidence from MetaSplice scores. (A) Likelihood ratios of the MetaSplice score intervals derived in the training calibration cohort, estimated in the 37,841 variants with at least one ClinVar review star in the test set’s calibration cohort (578 P/LP), with exact 95% CIs. Vertical bars mark the requirement for each level at a prior of 0.0441. The candidate rule combines the PP3 Strong and Very Strong intervals, and the BP4 Moderate and Strong intervals, at Supporting strength. (B) Share of variants given PP3, BP4 or no code by the candidate MetaSplice rule (PP3 for scores ≥ 0.544, BP4 for scores ≤ 0.0017, both at Supporting strength) and by the ClinGen SpliceAI rule (PP3 for scores ≥ 0.2, BP4 for scores ≤ 0.1, using SpliceAI scores without masking over ±4,999 nucleotides), in the test set’s calibration cohort and among 859 resolved variants of uncertain significance. Variants without a score receive no code. CI, confidence interval; P/LP, pathogenic or likely pathogenic; B/LB, benign or likely benign; VUS, variants of uncertain significance.

Because some intervals were inconclusive and the prior was carried over from missense calibration, we defined a candidate rule that uses MetaSplice at Supporting strength only. The rule’s PP3 group comprises the PP3 Strong and Very Strong intervals, and its BP4 group the BP4 Moderate interval together with the dropped BP4 Strong interval (Supplemental Methods). In the test set’s calibration cohort, the rule gave PP3 to 505 of 578 P/LP variants (87.4%) and to 73 of 37,263 B/LB variants (0.20%), and BP4 to 32,548 B/LB variants (87.3%) and 12 P/LP variants (2.1%) (Figure 5B; Table 2). The ClinGen SpliceAI rule gave PP3 to 524 P/LP (90.7%) and 1,654 B/LB variants (4.44%), and BP4 to 32,966 B/LB (88.5%) and 39 P/LP variants (6.7%). MetaSplice thus gave fewer discordant codes: PP3 to 4.24 percentage points fewer B/LB variants (95% CI 4.03–4.46) and BP4 to 4.67 points fewer P/LP variants (2.77–6.57). It also gave fewer concordant codes: PP3 to 3.29 points fewer P/LP variants (0.87–5.88, or −0.98 to 6.94 with genes resampled) and BP4 to 1.12 points fewer B/LB variants (0.74–1.49). More variants received no code (10.6% of P/LP and 12.5% of B/LB variants, compared with 2.6% and 7.1%). The PP3 and BP4 groups of both rules met the Supporting requirement by CI at priors of 0.01, 0.0441 and 0.1, with PP3 likelihood ratios of 446 (95% CI 331–609) for MetaSplice and 20.4 (18.7–22.2) for SpliceAI. SpliceAI’s precomputed masked scores and the maximum over all genes gave the same direction of difference for PP3 among B/LB variants (Table S9).

**Table 2.** Codes assigned by the candidate MetaSplice rule and the ClinGen SpliceAI rule. Variants with at least one ClinVar review star in the test set’s calibration cohort, and resolved variants of uncertain significance (uncertain or conflicting in the training release, P/LP or B/LB in September 2026).

| Set | Rule | P/LP with PP3 | B/LB with PP3 | B/LB with BP4 | P/LP with BP4 | P/LP, no code | B/LB, no code | PP3 likelihood ratio [95% CI] | BP4 likelihood ratio [95% CI] |
| --- | --- | --- | --- | --- | --- | --- | --- | --- | --- |
| Calibration cohort (578 P/LP; 37,263 B/LB) | MetaSplice ($\geq 0.544 / \leq 0.0017$)<sup>a</sup> | 505 (87.4%) | 73 (0.20%) | 32,548 (87.3%) | 12 (2.1%) | 61 (10.6%) | 4,642 (12.5%) | 446 [331, 609] | 0.0238 [0.0111, 0.0441] |
| Calibration cohort (578 P/LP; 37,263 B/LB) | SpliceAI ($\geq 0.2 / \leq 0.1$)<sup>b</sup> | 524 (90.7%) | 1,654 (4.44%) | 32,966 (88.5%) | 39 (6.7%) | 15 (2.6%) | 2,643 (7.1%) | 20.4 [18.7, 22.2] | 0.0763 [0.0519, 0.107] |
| Resolved VUS (153 P/LP; 706 B/LB) | MetaSplice ($\geq 0.544 / \leq 0.0017$)<sup>a</sup> | 139 (90.8%) | 7 (0.99%) | 512 (72.5%) | 0 (0.0%) | 14 (9.2%) | 187 (26.5%) | 91.6 [38.3, 276] | 0 [0, 0.0412] |
| Resolved VUS (153 P/LP; 706 B/LB) | SpliceAI ($\geq 0.2 / \leq 0.1$)<sup>b</sup> | 145 (94.8%) | 220 (31.16%) | 455 (64.4%) | 8 (5.2%) | 0 (0.0%) | 31 (4.4%) | 3.04 [2.53, 3.59] | 0.0811 [0.0293, 0.179] |

<sup>a</sup>Both codes at Supporting strength; thresholds rounded (exact values in Table S3). <sup>b</sup>ClinGen thresholds, applied at Supporting strength to SpliceAI 1.3.1 scores without masking over $\pm 4,999$ nucleotides. A variant without a score receives no code. Likelihood-ratio CIs are exact, combining 97.5% Clopper–Pearson intervals for the two class proportions. Results for all review statuses and for other SpliceAI configurations are given in Table S9. B/LB, benign or likely benign; CI, confidence interval; P/LP, pathogenic or likely pathogenic; VUS, variant of uncertain significance.

Among 859 variants that were uncertain or conflicting in the training release and have since been classified (153 P/LP, 706 B/LB), MetaSplice gave PP3 to 7 later-B/LB variants (0.99%), compared with 220 (31.2%) under the SpliceAI rule (difference 30.2 percentage points, 95% CI 26.8–33.6), and BP4 to none of the later-P/LP variants, compared with 8 (5.2%). It gave PP3 to 139 later-P/LP variants (90.8%), compared with 145 (94.8%) under the SpliceAI rule, and BP4 to 512 later-B/LB variants (72.5%), compared with 455 (64.4%). In this subgroup, the SpliceAI rule’s PP3 likelihood ratio was 3.04 (2.53–3.59), which met the Supporting requirement by CI at priors of 0.0441 and 0.1 but not at 0.01. MetaSplice’s was 91.6 (38.3–276). Among 60,776 VUS that remained unresolved, MetaSplice gave PP3 to 14,939 (24.6%), BP4 to 21,482 (35.3%) and neither to 24,355 (40.1%). The SpliceAI rule gave PP3 to 23,508 (38.7%), BP4 to 31,179 (51.3%) and neither to 6,089 (10.0%) (Table S10).

## Discussion

MetaSplice separates P/LP from B/LB intronic variants outside the canonical splice sites better than SpliceAI and CADD, as measured by auPRC, with the largest gain at deep intronic positions, where SpliceAI’s auPRC is lowest. The gain did not require the variant’s gene to have been seen in training, and most of it remained after the gene-level features or DANN and GPN-MSA were removed. A candidate PP3/BP4 rule derived from its ClinVar calibration gave pathogenic-supporting evidence to far fewer benign test variants, and benign-supporting evidence to fewer pathogenic ones, than the ClinGen SpliceAI rule.

The comparison with its own inputs shows where the ensemble helps. Pangolin alone ranked non-canonical variants almost as well as the splicing-only baselines, and MetaSplice’s gain over it was concentrated at deep intronic positions and in the donor region. In the acceptor and PPT regions, it was not clear. Within the genes that contain both P/LP and B/LB variants, MetaSplice and SpliceAI ranked variants similarly. That analysis uses a different measure and a subset of genes, so it does not resolve whether the pooled advantage arises mainly within or between genes. The gene-disjoint analysis shows that the advantage does not require the variant’s gene to have been seen in training. Like our earlier dbscSNV,^9^ MetaSplice combines complementary predictors, but it scores every intronic position of MANE transcripts and is trained on clinical classifications.

MetaSplice predicts clinical pathogenicity, not splicing outcome. Against experimentally assayed splicing, it exceeded SpliceAI in the donor and acceptor regions but did not clearly differ in the PPT, branch-point and deep intronic regions. When low-frequency splice-altering variants were added to the positives, within-region AUROC fell to near 0.5 for all three tools in those regions. That combined endpoint does not test low-frequency variants against normal variants separately, and because low-frequency aberrant transcripts need not reduce gene function, it does not by itself limit a pathogenicity predictor. SpliceAI, SpliceTransformer and Pangolin remain the tools of choice when the question is the molecular consequence of a variant. MetaSplice complements them when the question is pathogenicity.

The candidate rule applies MetaSplice at Supporting strength only, at its PP3 Strong and BP4 Moderate thresholds. It would replace rather than add to a SpliceAI-based PP3 or BP4, and observed-splicing codes would take precedence when RNA assays are available.^19^ In this test set, it assigned fewer codes discordant with ClinVar classifications than the ClinGen SpliceAI rule but left more variants without a code (40.1% of remaining VUS, compared with 10.0%). The rule was chosen after test-set results had been examined, and it rests on a prior carried over from missense calibration. It is therefore a candidate for independent evaluation, including its use at Supporting strength, in variants classified after this study and with a prior estimated for intronic variants. It is not yet a recommendation for clinical use.

Several limitations affect interpretation. First, ClinVar classifications can rest partly on computational evidence, including SpliceAI, and ClinVar variants are not a random sample of intronic variants. Second, besides the choice of the rule, a candidate BP4 Strong level was dropped after likelihood ratios in test variants had been examined (Supplemental Methods), so the rule’s test-set results are a reassessment, not an independent validation. Third, some regions contain few P/LP variants: the branch-point region could not be evaluated, and in the PPT region MetaSplice and SpliceAI did not clearly differ. Finally, the scope excludes exonic variants and transcripts outside MANE, SpliceVarDB combines assays with different detection criteria, and the literature set contains positives only.

MetaSplice adds conservation, exon context and gene-level information to splicing predictions to rank intronic variants outside the canonical sites by pathogenicity, and its calibration yields a candidate PP3/BP4 rule for independent evaluation.

## Acknowledgments

This work was supported by the University of South Florida Provost’s Collaborative Research Excellence and Translational Efforts (CREATE) Award. The funder had no role in the design of the study, in the collection, analysis or interpretation of data, in the decision to publish or in the preparation of the manuscript. The author thanks the ClinVar, GENCODE, gnomAD and UCSC Genome Browser teams and the developers of the input prediction tools for making their resources publicly available.

## Author contributions

X.L. conceived and designed the study, curated the data, developed the software, performed all analyses and wrote the manuscript.

## Web resources

CADD, https://cadd.bihealth.org/

ClinGen Dosage Sensitivity, https://search.clinicalgenome.org/kb/gene-dosage

ClinVar, https://www.ncbi.nlm.nih.gov/clinvar/

dbNSFP (version 5.4), https://www.dbnsfp.org/

Ensembl assembly mapping (GRCh37–GRCh38 chain), https://ftp.ensembl.org/pub/assembly_mapping/homo_sapiens/

GENCODE, https://www.gencodegenes.org/ gnomAD, https://gnomad.broadinstitute.org/

GPN-MSA GRCh38 scores, https://huggingface.co/datasets/songlab/gpn-msa-hg38-scores HGNC, https://www.genenames.org/

MANE, https://www.ncbi.nlm.nih.gov/refseq/MANE/ MetaSplice, https://github.com/xiaoming-liu/MetaSplice

MetaSplice annotation data (mirror), https://usf.box.com/s/44v8lpk6rxi8d6vdvw1032cfr5olmzdq

MetaSplice Docker image, https://github.com/xiaoming-liu/MetaSplice/pkgs/container/metasplice

MetaSplice reproducibility package, https://github.com/xiaoming-liu/MetaSplice-reproducibility Pangolin, https://github.com/tkzeng/Pangolin

SpliceAI, https://github.com/Illumina/SpliceAI

SpliceTransformer, https://github.com/ShenLab-Genomics/SpliceTransformer SpliceVarDB, https://splicevardb.org/

UCSC Genome Browser downloads (phyloP and phastCons, GRCh38), https://hgdownload.soe.ucsc.edu/goldenPath/hg38/

XGBoost, https://xgboost.readthedocs.io/

## Data and code availability

The MetaSplice 2.0 code and model are available at https://github.com/xiaoming-liu/MetaSplice (release v2.0.1) and as the Docker image ghcr.io/xiaoming-liu/metasplice:2.0.1 (digest sha256:c32fc8d49da016efd39bc0ac3738ee46d873476e775c0a01a6c714621914f105), which scores variants from a VCF file. The annotation data it reads (219 GB) are mirrored at https://usf.box.com/s/44v8lpk6rxi8d6vdvw1032cfr5olmzdq. The reproducibility package, with the variant sets, models, analysis scripts and results of this study, is available at https://github.com/xiaoming-liu/MetaSplice-reproducibility (release v2.0.1). SpliceAI and CADD scores and the splicing classifications of the SpliceVarDB set are not redistributed, because their providers’ terms do not grant redistribution; the package includes scripts that rebuild them from the original sources. MetaSplice is licensed for non-commercial research, educational and clinical (non-revenue-generating) use; for commercial use, including internal use by for-profit entities, contact the author. This study analyzes publicly available data from ClinVar, gnomAD, GENCODE and the other resources listed under web resources.

### Declaration of generative AI and AI-assisted technologies in the manuscript preparation process

During the preparation of this work the author used Claude Opus 4.6 and Claude Opus 5 (Anthropic), GPT-6 (Astra), Codex and Claude Opus 5.5 in Claude Code (Anthropic) in order to draft, edit and reformat manuscript text. After using these tools, the author reviewed and edited the content as needed and takes full responsibility for the content of the publication.

