## Supplemental Methods and Tables for "MetaSplice: pathogenicity prediction and ACMG/AMP evidence for intronic variants outside the canonical splice sites"

**Variant processing**

ClinVar variants were read from the GRCh38 VCF releases of 31 August 2025, 15 April 2026 and 28 September 2026. We kept single-nucleotide variants (SNVs) and labeled them from the clinical significance field: pathogenic, likely pathogenic or pathogenic/likely pathogenic as P/LP, and benign, likely benign or benign/likely benign as B/LB. Records were deduplicated by chromosome, position, reference and alternate allele; a variant with conflicting labels across records would have been dropped, but none occurred.

Regions, genes and exon context were taken from GENCODE v50 (Ensembl 116) transcripts tagged MANE Select or MANE Plus Clinical: 19,373 transcripts, 19,299 genes and 204,067 exons. Each gene record was reset to the extent of its MANE transcripts; this changed 17,072 of the 19,299 records, whose GENCODE span also covered other transcripts. A variant was in scope if it lay within the span of a MANE transcript of its gene and outside every MANE exon, of any gene; the region rule is given in the main text and Table 1. Where a position belonged to several genes, assignment followed the region priority, then the nearest exon boundary, with remaining ties resolved by the fixed order of the annotation. Deep intronic positions inside an exon of any GENCODE v50 basic transcript were out of scope. In the training release, 1,370,180 SNVs were classified P/LP or B/LB; 413,199 were in scope, and the others were in a MANE exon (921,807), inside a GENCODE basic exon (22,569), outside MANE transcripts (10,851) or on a chromosome without a MANE transcript (1,754).

The temporal test set was drawn from the 457,664 in-scope SNVs classified P/LP or B/LB on 28 September 2026. The 411,552 present in the training data, and any other variant classified P/LP or B/LB in the training release (matched by allele or ClinVar identifier in either genome build), were removed, leaving 46,112. Labels, clinical significance and review status came from the 28 September 2026 release. Every test variant had exactly one feature row with the same region, gene and gene identifier as the variant list.

Variants of uncertain significance (VUS) were the 71,267 in-scope SNVs classified uncertain (56,282) or conflicting (14,985) on 15 April 2026 and absent from the training data. Of these, 61,016 met the calibration-cohort definition, and 240 of those had been classified P/LP or B/LB by 28 September 2026, leaving 60,776 remaining VUS. Resolved VUS were drawn from the test set's calibration cohort: variants uncertain or conflicting in the training release.

Literature-confirmed variants were extracted from PubMed abstracts describing experimentally supported intronic splice-altering variants, using Claude Opus 4.6 (Anthropic). A blinded manual review of 50 randomly selected candidates found 49 fully and one partially supported. The 746 records were lifted from GRCh37 to GRCh38 with the Ensembl assembly chain, requiring a one-to-one position and an identical reference allele and complementing both alleles where the mapped segment was inverted. One record failed (reference allele differs between builds); 727 were in scope (11 lay inside a GENCODE basic exon and 7 outside MANE transcripts); 30 were training variants, leaving 697.

The SpliceVarDB set was rebuilt from SpliceVarDB's source studies. We applied the criteria in SpliceVarDB's Table 1 to each study's published data. For SAVNet (Shiraishi et al. 2018) these are supporting read counts and Bayes factors (its Supplemental Table S3); a variant whose instances were classified differently was excluded. For MFASS (Chong et al. 2019) we used the change in exon inclusion index from the authors' processed data (v2_dpsi, or v1_dpsi where v2_dpsi was missing). For MiSplice (Jayasinghe et al. 2018) we used every call that passed manual review (its supplementary file mmc2), all of which are splice-altering. The rebuilt sets contained 13,925 SAVNet, 28,854 MFASS and 1,914 MiSplice variants; SpliceVarDB lists 13,864, 28,962 and 559. Class counts also differed: for MFASS, 1,256 splice-altering variants, compared with SpliceVarDB's 1,468. MaPSY was not included because its classifications could not be reproduced. Four literature studies with variant-level data were added: Rhine et al. 2022, Casadei et al. 2019, Fraile-Bethencourt et al. 2019 and Valenzuela-Palomo et al. 2022. A fifth, Sakaguchi and Suyama 2021, was excluded because its alternative alleles were not reported.

SpliceVarDB records (44,510) were lifted in the same way: 44,324 were lifted, 138 had no one-to-one position and 48 a different reference allele. Of the lifted records, 24,372 were in scope (19,375 were in a MANE exon, 431 inside a GENCODE basic exon and 146 outside MANE transcripts), and 1,464 were training variants, leaving 22,908 records for 22,314 unique SNVs (594 variants were assayed in two studies). Annotations were shared across records of the same variant. Forty variants called splice-altering by one assay and low-frequency splice-altering by another were excluded from the primary analysis, leaving 22,274 (8,015 splice-altering, 9,543 low-frequency splice-altering and 4,716 normal). The strict endpoint compared 8,015 splice-altering with 4,716 normal variants; the broad endpoint counted low-frequency splice-altering variants as positives.

**Features**

All coordinate-based lookups used GRCh38, except DANN, which is distributed for GRCh37 and was read at positions mapped back with the Ensembl chain after a reference-allele check. Conservation scores were phyloP and phastCons from the UCSC 100-way vertebrate and 470-way mammalian alignments. GPN-MSA scores were the published GRCh38 scores. Gene-level annotations (pLI, LOEUF and the ClinGen haploinsufficiency score) came from the dbNSFP 5.4 gene file, matched by Ensembl gene identifier (19,230 of 19,299 MANE genes found); values unavailable for a gene were left missing. The same values are obtained by rebuilding them from gnomAD, ClinGen and HGNC with the same rules. SpliceTransformer provided its aggregate score and 15 tissue scores; its gene annotation was rebuilt from the MANE transcripts, and GPU runs disabled TensorFloat-32 arithmetic. Pangolin provided the largest predicted increase, the largest predicted decrease and the larger of their absolute values, using a database built from the same MANE annotation. Exon context comprised the length of the nearest exon of the assigned gene, that length modulo three and the distance to its boundary. Scores were joined to variants by allele key. The 38 features are listed in Table S2. Population allele frequency was recorded for each variant to define the calibration cohort but was not a model input.

Before training or evaluation, region, gene and gene identifier were checked for agreement between each variant list and its feature table, and gene-level values were required to be identical for the same gene across datasets. As an end-to-end check, the command-line scorer was rerun from the input variants, including fresh SpliceTransformer and Pangolin runs, for 1,000 training and 986 test variants (up to 200 per non-canonical region). Region, gene and gene identifier matched for every variant; 37 of the 38 features were identical bit for bit, and the remaining feature (pLI) differed only in text representation. Scores from the command-line scorer, then written to six decimals, agreed with those from the stored features to that precision (largest difference 5.0 × 10^−7^), and fold-model predictions reproduced the out-of-fold scores to within 1.1 × 10^−16^. The MetaSplice 2.0 Docker image writes full-precision scores; run from the input variants on a GPU, it reproduced the published scores of 2,200 ClinVar variants (200 per non-canonical region and 50 per canonical site from each of the training and test sets) bit for bit.

**Comparator scores**

Precomputed SpliceAI scores were the masked GRCh37 SNV files (version 1.3), read at the GRCh37 position of each variant; the score was the largest of the four delta scores in the first record for the allele. CADD v1.7 PHRED-scaled scores were read for GRCh38. For the comparison with the ClinGen SpliceAI rule, SpliceAI 1.3.1 (TensorFlow 2.15.1) was run without masking over ±4,999 nucleotides (options -A grch37 -D 4999 -M 0), the configuration in which ClinGen calibrated its thresholds. Before scoring, the installation was checked by reproducing the precomputed masked scores (options -D 50 -M 1) for 500 variants (up to 100 per non-canonical region): 99.20% were within 0.01. The primary value was the largest delta score of the first record for the allele; the largest value over all overlapping genes was a sensitivity analysis. Raw and precomputed scores covered the same variants in every cohort.

**Model training and evaluation**

Inputs were converted to 32-bit floating-point values. For each fit, the positive-class weight was the ratio of B/LB to P/LP variants in that fit. For gene-grouped cross-validation, Ensembl gene identifiers were shuffled with seed 42 and assigned to five folds. The final model was trained on all training variants. For the gene-disjoint analysis, the 7,194 genes of the training and test sets were assigned to five folds with seed 42; each fold model was trained on training variants of the other folds and scored the test variants of its fold, so every test variant was scored by a model that had not seen its gene.

auPRC was computed as average precision. Single-tool CIs came from 1,000 bootstrap resamples drawn separately within the P/LP and B/LB classes. Paired differences used the same resampling, recalculating both tools' auPRC on each resample of the variants both tools score. For gene resampling, genes were drawn with replacement and each variant weighted by the number of times its gene was drawn (1,000 resamples, seed 42). Within-gene concordance was the proportion of P/LP–B/LB pairs within the same gene in which the P/LP variant scored higher, with ties counted as one half, pooled over genes with both classes. SHAP values were computed on the model's log-odds scale.

**Calibration**

Likelihood ratios were estimated following Pejaver et al. 2022. Scores were evaluated at every distinct score when there were at most 2,000 (101 for the precomputed SpliceAI scores) and otherwise on a grid combining 1,501 quantiles of the score distribution with 501 evenly spaced scores between its minimum and maximum (2,000 distinct points for MetaSplice). At each grid score, a symmetric window was widened until it held at least 100 labeled variants and at least 25 of each class, and the local likelihood ratio was the share of P/LP variants in the window divided by the share of B/LB variants. Uncertainty came from 10,000 bootstrap resamples drawn within each class. A PP3 level was reached at the smallest grid score at and above which the 5th percentile of the bootstrap likelihood ratio met the level's requirement everywhere; a BP4 level at the largest grid score at and below which the 95th percentile met the reciprocal requirement everywhere. The requirements follow from a prior probability of pathogenicity of 0.0441 (c = 1,124): 2.41, 5.79, 33.5 and 1,124 for Supporting, Moderate, Strong and Very Strong. Thresholds were derived separately for variants with at least one review star (the main analysis) and for all review statuses (Table S3).

To estimate a prior for rare non-canonical intronic variants, we scored 100,000 variants sampled uniformly from the eligible gnomAD population (and 50,000 stratified by region), each with the fold model that excluded its gene, and fitted the reference score distribution as a mixture of the out-of-fold P/LP and B/LB score distributions on 50 bins of the logit score. The pre-set criteria for adopting the estimate were a cumulative-distribution discrepancy of at most 0.02, estimates with 25 and 100 bins within a factor of 1.5 of the primary estimate, and a CI excluding 0.0441. The primary estimate was 0.0311 (95% CI 0.0286–0.0340), but the discrepancy was 0.41 and the 25- and 100-bin estimates were 0.084 and 0.0065, so 0.0441 was retained.

In the test set, each interval's likelihood ratio was its share of P/LP variants divided by its share of B/LB variants. Exact 95% CIs combined 97.5% Clopper–Pearson intervals for the two shares, which keeps a defined bound when an interval contains no variants of one class. A level's CI cleared the requirement when its lower bound (PP3) or upper bound (BP4) met it.

**Pre-specification**

Each analysis was written into a dated protocol, with the checksums of its input files, before it was computed. Before computing any performance measure, the scripts of the test-set and VUS analyses verified those checksums and the expected denominators; the scripts deriving the thresholds for variants with at least one review star and those of the SpliceVarDB and literature analyses verified the expected denominators. The model was trained once and was not changed after test-set evaluation began. Two decisions about thresholds used test variants. The candidate BP4 Strong level was dropped after its likelihood ratio in an earlier evaluation of test variants did not meet the Strong requirement, and the Supporting-only rule was chosen after interval likelihood ratios in test variants had been examined. The scripts and their logs are part of the analysis package.

**Supplemental tables**

**Table S1. Datasets other than the ClinVar training and test sets**

| Dataset | Variants | Use |
| --- | --- | --- |
| SpliceVarDB, after removal of training variants | 22,274 (17,558 splice-altering, including low-frequency; 4,716 normal) | Measured splicing. The strict endpoint uses 12,731: 8,015 splice-altering and 4,716 normal. |
| Literature-confirmed splice-altering variants | 697, all P/LP, none in training | Recall at fixed scores (positives only). |
| Resolved VUS: uncertain or conflicting in ClinVar 2025-08-31, P/LP or B/LB in 2026-09-28 | 859 at ≥ 1 review star (153 P/LP, 706 B/LB), the set reported in the main text; 861 at all review statuses (Table S9b) | Codes under the candidate PP3/BP4 rule and under ClinGen's SpliceAI rule. A subgroup of the test set's calibration cohort. |
| Remaining VUS: uncertain or conflicting in ClinVar 2026-04-15, not P/LP or B/LB in 2026-09-28 | 60,776 in the calibration cohort | Score distribution, and codes under each rule. |
| gnomAD v4.1 reference sample | 100,000 uniform and 50,000 stratified by region | Attempted estimate of the prior (Supplemental methods) |

B/LB, benign or likely benign; P/LP, pathogenic or likely pathogenic; VUS, variant of uncertain significance.

**Table S2. The 38 MetaSplice features**

| Group | Features | Number | Source |
| --- | --- | --- | --- |
| SpliceTransformer | Aggregate score and 15 tissue scores (adipose tissue, blood, blood vessel, brain, colon, heart, kidney, liver, lung, muscle, nerve, small intestine, skin, spleen, stomach) | 16 | SpliceTransformer, GPU, MANE annotation |
| Pangolin | Largest predicted increase, largest predicted decrease, larger absolute value | 3 | Pangolin, MANE database |
| Conservation | phyloP and phastCons, 100-way vertebrate and 470-way mammalian | 4 | UCSC, GRCh38 |
| GPN-MSA | Variant score | 1 | Published GRCh38 scores |
| DANN | Variant score | 1 | DANN, GRCh37 position |
| Gene-level | pLI, LOEUF, ClinGen haploinsufficiency score | 3 | dbNSFP 5.4 gene file; rebuilt from gnomAD, ClinGen and HGNC with the same rules |
| Exon context | Nearest exon length, length modulo 3, distance to the exon boundary | 3 | GENCODE v50 MANE |
| Region | Indicators for the seven regions | 7 | GENCODE v50 MANE |

The tissue list follows SpliceTransformer's 15 tissue outputs; the model order of all 38 features is given in the reproducibility package (data and code availability). LOEUF, loss-of-function observed/expected upper bound fraction; pLI, probability of loss-of-function intolerance.

**Table S3. Score thresholds derived in the training calibration cohort**

Thresholds at which out-of-fold scores met each likelihood-ratio requirement at a prior of 0.0441, for variants with at least one ClinVar review star (main analysis) and for all review statuses.

| Tool | Level | Likelihood ratio required | At least one star | All review statuses |
| --- | --- | --- | --- | --- |
| MetaSplice | PP3 Very Strong | 1124.0000 | ≥ 0.993946915744262 | ≥ 0.9937176871299744 |
| MetaSplice | PP3 Strong | 33.5261 | ≥ 0.5439710286928215 | ≥ 0.547970814355501 |
| MetaSplice | PP3 Moderate | 5.7902 | ≥ 0.1399926767621949 | ≥ 0.1579917122442525 |
| MetaSplice | PP3 Supporting | 2.4063 | ≥ 0.0519973921832465 | ≥ 0.0499974993519067 |
| MetaSplice | BP4 Supporting | 0.4156 | ≤ 0.0038620028824855 | ≤ 0.0028122577661026 |
| MetaSplice | BP4 Moderate | 0.1727 | ≤ 0.0017023810017853 | ≤ 0.001078249971227 |
| MetaSplice | BP4 Strong | 0.0298 | ≤ 0.0002745522665208 | ≤ 0.0001829224231186 |
| SpliceAI | PP3 Very Strong | 1124.0000 | not reached | not reached |
| SpliceAI | PP3 Strong | 33.5261 | ≥ 0.49 | ≥ 0.49 |
| SpliceAI | PP3 Moderate | 5.7902 | ≥ 0.3 | ≥ 0.3 |
| SpliceAI | PP3 Supporting | 2.4063 | ≥ 0.2 | ≥ 0.2 |
| SpliceAI | BP4 Supporting | 0.4156 | ≤ 0.08 | ≤ 0.04 |
| SpliceAI | BP4 Moderate | 0.1727 | ≤ 0.03 | ≤ 0.01 |
| SpliceAI | BP4 Strong | 0.0298 | not reached | not reached |

The candidate rule uses the at-least-one-star MetaSplice thresholds for PP3 Strong (0.5439710286928215) and BP4 Moderate (0.0017023810017853637) at Supporting strength; the main text rounds them to 0.544 and 0.0017. Rounding the BP4 threshold to 0.0017 would change 9 BP4 assignments in the test set's calibration cohort. The BP4 Strong level was not selected. SpliceAI thresholds were derived from precomputed masked scores.

**Table S4. Gene-grouped cross-validation in the training set**

Out-of-fold predictions from five gene-grouped folds.

| Set | Variants | P/LP | auPRC | AUROC |
| --- | --- | --- | --- | --- |
| All training variants | 413,199 | 41,032 | 0.9942 | 0.9987 |
| Non-canonical | 375,497 | 3,599 | 0.8779 | 0.9870 |
| Canonical donor site | 20,576 | 20,443 | 0.9983 | 0.8289 |
| Canonical acceptor site | 17,126 | 16,990 | 0.9972 | 0.8063 |
| Donor region | 26,476 | 1,862 | 0.9559 | 0.9953 |
| Acceptor region | 64,628 | 784 | 0.8562 | 0.9797 |
| PPT region | 86,776 | 348 | 0.7105 | 0.9757 |
| Branch-point region | 5,822 | 53 | 0.5551 | 0.9325 |
| Deep intronic | 191,795 | 552 | 0.7132 | 0.9695 |
| Deep intronic, sensitivity set | 159,226 | 364 | 0.7325 | 0.9646 |

auPRC, area under the precision–recall curve; AUROC, area under the receiver operating characteristic curve; B/LB, benign or likely benign; CI, confidence interval; P/LP, pathogenic or likely pathogenic; PPT, polypyrimidine tract.

**Table S5. Discrimination on the temporal test set**

Each tool's auPRC on the variants it scores; differences on the variants both tools score, with CIs from class-stratified resampling and then from gene resampling.

| Region | Variants (P/LP) | MetaSplice | SpliceAI | CADD | MetaSplice, SpliceAI-scored (n) | MetaSplice minus SpliceAI | MetaSplice minus CADD |
| --- | --- | --- | --- | --- | --- | --- | --- |
| Non-canonical | 40,619 (602) | 0.9160 | 0.7566 | 0.3638 | 0.9188 (40,296) | +0.1622 [0.1336, 0.1923]; [0.1284, 0.1971] | +0.5522 [0.5117, 0.5899]; [0.5102, 0.5952] |
| Donor region | 3,314 (283) | 0.9646 | 0.8646 | 0.6878 | 0.9693 (3,278) | +0.1047 [0.0738, 0.1378]; [0.0669, 0.1476] | +0.2768 [0.2293, 0.3272]; [0.2168, 0.3409] |
| Acceptor region | 7,021 (123) | 0.9468 | 0.8645 | 0.3486 | 0.9468 (6,983) | +0.0823 [0.0362, 0.1292]; [0.0359, 0.1370] | +0.5982 [0.5110, 0.6792]; [0.5059, 0.6850] |
| PPT region | 12,410 (61) | 0.7825 | 0.7283 | 0.1090 | 0.7825 (12,310) | +0.0542 [−0.0245, 0.1356]; [−0.0290, 0.1377] | +0.6735 [0.5473, 0.7691]; [0.5522, 0.7662] |
| Branch-point region | 395 (9) | counts only |  |  |  |  |  |
| Deep intronic | 17,479 (126) | 0.8241 | 0.5510 | 0.3275 | 0.8241 (17,331) | +0.2731 [0.2046, 0.3419]; [0.2038, 0.3575] | +0.4966 [0.4141, 0.5761]; [0.4243, 0.5701] |
| Deep intronic, sensitivity set | 14,459 (99) | 0.8468 | 0.6070 | 0.3523 | 0.8468 (14,325) | +0.2398 [0.1688, 0.3110]; [0.1664, 0.3285] | +0.4945 [0.4016, 0.5902]; [0.4115, 0.5767] |
| Canonical sites | 5,493 (5,454) | 0.9975 | 0.9946 | 0.9950 | 0.9975 (5,453) | +0.0029 [0.0012, 0.0045]; [0.0008, 0.0053] | +0.0025 [0.0010, 0.0039]; [0.0002, 0.0056] |
| All test variants | 46,112 (6,056) | 0.9947 | 0.9763 | 0.9707 | 0.9948 (45,749) | +0.0185 [0.0151, 0.0215]; [0.0137, 0.0243] | +0.0241 [0.0212, 0.0271]; [0.0203, 0.0287] |

Regions with fewer than 20 P/LP variants are counts only. MetaSplice and CADD score every test variant. auPRC, area under the precision–recall curve; AUROC, area under the receiver operating characteristic curve; B/LB, benign or likely benign; CI, confidence interval; P/LP, pathogenic or likely pathogenic; PPT, polypyrimidine tract.

**Table S6. Single inputs and baselines on non-canonical test variants**

Each input or baseline on the variants it scores; MetaSplice minus each on the variants both score.

| Region | Input or baseline | Coverage | auPRC | AUROC | MetaSplice minus [95% CI] |
| --- | --- | --- | --- | --- | --- |
| Non-canonical | Pangolin max | 100.0% | 0.8563 | 0.9764 | +0.0597 [0.0394, 0.0803] |
| Non-canonical | Pangolin increase | 100.0% | 0.4844 | 0.8985 | +0.4315 [0.3922, 0.4687] |
| Non-canonical | Pangolin decrease (negated) | 100.0% | 0.7240 | 0.8571 | +0.1920 [0.1607, 0.2208] |
| Non-canonical | SpliceTransformer | 100.0% | 0.7169 | 0.9594 | +0.1991 [0.1671, 0.2279] |
| Non-canonical | GPN-MSA (negated) | 100.0% | 0.2809 | 0.7784 | +0.6350 [0.5953, 0.6737] |
| Non-canonical | DANN | 99.9% | 0.1060 | 0.7437 | +0.8099 [0.7840, 0.8304] |
| Non-canonical | phyloP 100-way | 100.0% | 0.2046 | 0.7831 | +0.7113 [0.6729, 0.7427] |
| Non-canonical | phyloP 470-way | 92.3% | 0.1730 | 0.7816 | +0.7407 [0.7059, 0.7716] |
| Non-canonical | phastCons 100-way | 100.0% | 0.0984 | 0.7852 | +0.8175 [0.7974, 0.8363] |
| Non-canonical | phastCons 470-way | 99.1% | 0.0769 | 0.7788 | +0.8384 [0.8203, 0.8577] |
| Non-canonical | pLI | 96.5% | 0.0186 | 0.5406 | +0.8981 [0.8795, 0.9172] |
| Non-canonical | LOEUF (negated) | 90.0% | 0.0196 | 0.5096 | +0.8858 [0.8649, 0.9077] |
| Non-canonical | XGBoost, 19 deep-learning features | 100.0% | 0.8794 | 0.9856 | +0.0366 [0.0203, 0.0529] |
| Non-canonical | logistic regression, 3 features | 100.0% | 0.8549 | 0.9789 | +0.0611 [0.0449, 0.0762] |
| Donor region | Pangolin max | 100.0% | 0.9390 | 0.9927 | +0.0255 [0.0130, 0.0392] |
| Donor region | Pangolin increase | 100.0% | 0.4594 | 0.8615 | +0.5051 [0.4480, 0.5589] |
| Donor region | Pangolin decrease (negated) | 100.0% | 0.9469 | 0.9924 | +0.0177 [0.0079, 0.0277] |
| Donor region | SpliceTransformer | 100.0% | 0.8327 | 0.9560 | +0.1318 [0.0981, 0.1678] |
| Donor region | GPN-MSA (negated) | 100.0% | 0.6183 | 0.9139 | +0.3463 [0.2877, 0.3976] |
| Donor region | DANN | 99.9% | 0.4924 | 0.8889 | +0.4721 [0.4082, 0.5241] |
| Donor region | phyloP 100-way | 100.0% | 0.6791 | 0.9359 | +0.2854 [0.2363, 0.3342] |
| Donor region | phyloP 470-way | 92.8% | 0.6845 | 0.9368 | +0.2764 [0.2261, 0.3298] |
| Donor region | phastCons 100-way | 100.0% | 0.4767 | 0.9260 | +0.4878 [0.4506, 0.5205] |
| Donor region | phastCons 470-way | 99.4% | 0.4407 | 0.9200 | +0.5238 [0.4939, 0.5536] |
| Donor region | pLI | 96.5% | 0.1075 | 0.5575 | +0.8573 [0.8342, 0.8763] |
| Donor region | LOEUF (negated) | 89.7% | 0.1118 | 0.5362 | +0.8527 [0.8174, 0.8774] |
| Donor region | XGBoost, 19 deep-learning features | 100.0% | 0.9469 | 0.9913 | +0.0177 [0.0058, 0.0308] |
| Donor region | logistic regression, 3 features | 100.0% | 0.9342 | 0.9903 | +0.0304 [0.0127, 0.0491] |
| Acceptor region | Pangolin max | 100.0% | 0.9158 | 0.9903 | +0.0310 [−0.0019, 0.0735] |
| Acceptor region | Pangolin increase | 100.0% | 0.6237 | 0.9148 | +0.3231 [0.2478, 0.4065] |
| Acceptor region | Pangolin decrease (negated) | 100.0% | 0.9178 | 0.9900 | +0.0290 [0.0034, 0.0595] |
| Acceptor region | SpliceTransformer | 100.0% | 0.8387 | 0.9815 | +0.1081 [0.0578, 0.1621] |
| Acceptor region | GPN-MSA (negated) | 100.0% | 0.2025 | 0.7829 | +0.7443 [0.6695, 0.8092] |
| Acceptor region | DANN | 99.9% | 0.0284 | 0.5858 | +0.9184 [0.8813, 0.9463] |
| Acceptor region | phyloP 100-way | 100.0% | 0.0512 | 0.6797 | +0.8956 [0.8485, 0.9238] |
| Acceptor region | phyloP 470-way | 93.6% | 0.0477 | 0.6962 | +0.9008 [0.8620, 0.9298] |
| Acceptor region | phastCons 100-way | 100.0% | 0.0386 | 0.6763 | +0.9082 [0.8750, 0.9366] |
| Acceptor region | phastCons 470-way | 99.8% | 0.0347 | 0.6804 | +0.9121 [0.8775, 0.9424] |
| Acceptor region | pLI | 96.2% | 0.0229 | 0.5315 | +0.9248 [0.8863, 0.9522] |
| Acceptor region | LOEUF (negated) | 89.8% | 0.0179 | 0.4829 | +0.9178 [0.8779, 0.9541] |
| Acceptor region | XGBoost, 19 deep-learning features | 100.0% | 0.9314 | 0.9905 | +0.0154 [−0.0085, 0.0397] |
| Acceptor region | logistic regression, 3 features | 100.0% | 0.9242 | 0.9904 | +0.0226 [−0.0024, 0.0485] |
| PPT region | Pangolin max | 100.0% | 0.7740 | 0.9799 | +0.0085 [−0.0485, 0.0736] |
| PPT region | Pangolin increase | 100.0% | 0.5956 | 0.9628 | +0.1869 [0.0895, 0.2894] |
| PPT region | Pangolin decrease (negated) | 100.0% | 0.7730 | 0.9697 | +0.0095 [−0.0426, 0.0747] |
| PPT region | SpliceTransformer | 100.0% | 0.6700 | 0.9646 | +0.1125 [0.0428, 0.1924] |
| PPT region | GPN-MSA (negated) | 100.0% | 0.0093 | 0.5642 | +0.7732 [0.6702, 0.8603] |
| PPT region | DANN | 99.8% | 0.0074 | 0.5888 | +0.7751 [0.6817, 0.8677] |
| PPT region | phyloP 100-way | 100.0% | 0.0087 | 0.6203 | +0.7738 [0.6754, 0.8627] |
| PPT region | phyloP 470-way | 92.0% | 0.0099 | 0.6165 | +0.7689 [0.6696, 0.8545] |
| PPT region | phastCons 100-way | 100.0% | 0.0061 | 0.5644 | +0.7764 [0.6779, 0.8658] |
| PPT region | phastCons 470-way | 99.4% | 0.0074 | 0.6001 | +0.7751 [0.6752, 0.8598] |
| PPT region | pLI | 96.4% | 0.0051 | 0.5071 | +0.7927 [0.6968, 0.8777] |
| PPT region | LOEUF (negated) | 89.9% | 0.0050 | 0.4899 | +0.7606 [0.6473, 0.8593] |
| PPT region | XGBoost, 19 deep-learning features | 100.0% | 0.7464 | 0.9784 | +0.0361 [−0.0276, 0.1040] |
| PPT region | logistic regression, 3 features | 100.0% | 0.7239 | 0.9748 | +0.0586 [−0.0011, 0.1161] |
| Deep intronic | Pangolin max | 100.0% | 0.7039 | 0.9228 | +0.1201 [0.0727, 0.1742] |
| Deep intronic | Pangolin increase | 100.0% | 0.7119 | 0.9282 | +0.1122 [0.0658, 0.1672] |
| Deep intronic | Pangolin decrease (negated) | 100.0% | 0.0323 | 0.3910 | +0.7918 [0.7258, 0.8526] |
| Deep intronic | SpliceTransformer | 100.0% | 0.5755 | 0.9393 | +0.2485 [0.1805, 0.3202] |
| Deep intronic | GPN-MSA (negated) | 100.0% | 0.0073 | 0.4848 | +0.8168 [0.7540, 0.8747] |
| Deep intronic | DANN | 99.9% | 0.0088 | 0.5616 | +0.8153 [0.7527, 0.8695] |
| Deep intronic | phyloP 100-way | 100.0% | 0.0098 | 0.5781 | +0.8130 [0.7504, 0.8679] |
| Deep intronic | phyloP 470-way | 92.0% | 0.0096 | 0.5645 | +0.8195 [0.7560, 0.8761] |
| Deep intronic | phastCons 100-way | 100.0% | 0.0096 | 0.6021 | +0.8132 [0.7510, 0.8688] |
| Deep intronic | phastCons 470-way | 98.7% | 0.0085 | 0.5588 | +0.8089 [0.7461, 0.8675] |
| Deep intronic | pLI | 96.7% | 0.0094 | 0.5299 | +0.8099 [0.7478, 0.8681] |
| Deep intronic | LOEUF (negated) | 90.1% | 0.0154 | 0.4983 | +0.7732 [0.6913, 0.8447] |
| Deep intronic | XGBoost, 19 deep-learning features | 100.0% | 0.7708 | 0.9687 | +0.0533 [0.0137, 0.0978] |
| Deep intronic | logistic regression, 3 features | 100.0% | 0.6874 | 0.9446 | +0.1367 [0.0928, 0.1855] |
| Deep intronic, sensitivity set | Pangolin max | 100.0% | 0.7699 | 0.9337 | +0.0769 [0.0371, 0.1233] |
| Deep intronic, sensitivity set | Pangolin increase | 100.0% | 0.7765 | 0.9343 | +0.0703 [0.0316, 0.1162] |
| Deep intronic, sensitivity set | Pangolin decrease (negated) | 100.0% | 0.0398 | 0.3978 | +0.8071 [0.7353, 0.8801] |
| Deep intronic, sensitivity set | SpliceTransformer | 100.0% | 0.6410 | 0.9322 | +0.2058 [0.1366, 0.2767] |
| Deep intronic, sensitivity set | GPN-MSA (negated) | 100.0% | 0.0072 | 0.4852 | +0.8396 [0.7765, 0.9041] |
| Deep intronic, sensitivity set | DANN | 99.9% | 0.0076 | 0.5497 | +0.8392 [0.7782, 0.9014] |
| Deep intronic, sensitivity set | phyloP 100-way | 100.0% | 0.0100 | 0.5851 | +0.8350 [0.7618, 0.8920] |
| Deep intronic, sensitivity set | phyloP 470-way | 91.8% | 0.0090 | 0.5507 | +0.8438 [0.7835, 0.9037] |
| Deep intronic, sensitivity set | phastCons 100-way | 100.0% | 0.0094 | 0.6096 | +0.8357 [0.7635, 0.8943] |
| Deep intronic, sensitivity set | phastCons 470-way | 98.7% | 0.0079 | 0.5436 | +0.8346 [0.7679, 0.8968] |
| Deep intronic, sensitivity set | pLI | 96.7% | 0.0105 | 0.5520 | +0.8326 [0.7660, 0.8914] |
| Deep intronic, sensitivity set | LOEUF (negated) | 90.3% | 0.0199 | 0.5325 | +0.8059 [0.7288, 0.8812] |
| Deep intronic, sensitivity set | XGBoost, 19 deep-learning features | 100.0% | 0.8224 | 0.9661 | +0.0245 [−0.0074, 0.0593] |
| Deep intronic, sensitivity set | logistic regression, 3 features | 100.0% | 0.7414 | 0.9475 | +0.1054 [0.0617, 0.1505] |

Inputs where lower values indicate pathogenicity were negated. Baselines: XGBoost on the 19 SpliceTransformer and Pangolin features; logistic regression on the aggregate SpliceTransformer score, Pangolin maximum and phyloP (100-way). auPRC, area under the precision–recall curve; AUROC, area under the receiver operating characteristic curve; B/LB, benign or likely benign; CI, confidence interval; P/LP, pathogenic or likely pathogenic; PPT, polypyrimidine tract.

**Table S7. Ablation models on non-canonical test variants**

Each ablation model retrained with the full model's protocol; differences on the variants both models or tools score (SpliceAI comparisons on the variants SpliceAI scores).

| Region | Ablation | auPRC [95% CI] | Minus full model | Minus SpliceAI | Minus CADD |
| --- | --- | --- | --- | --- | --- |
| Non-canonical | (a) without gene-level | 0.9054 [0.8846, 0.9253] | −0.0105 [−0.0183, −0.0029] | +0.1518 [0.1227, 0.1826] | +0.5417 [0.5019, 0.5781] |
| Donor region | (a) without gene-level | 0.9581 [0.9375, 0.9754] | −0.0065 [−0.0153, 0.0012] | +0.0983 [0.0668, 0.1324] | +0.2703 [0.2242, 0.3179] |
| Acceptor region | (a) without gene-level | 0.9509 [0.9198, 0.9764] | +0.0041 [−0.0072, 0.0153] | +0.0864 [0.0433, 0.1337] | +0.6023 [0.5171, 0.6806] |
| PPT region | (a) without gene-level | 0.7557 [0.6546, 0.8535] | −0.0268 [−0.0656, 0.0083] | +0.0274 [−0.0555, 0.1256] | +0.6467 [0.5163, 0.7592] |
| Deep intronic | (a) without gene-level | 0.7858 [0.7123, 0.8536] | −0.0383 [−0.0738, −0.0076] | +0.2348 [0.1586, 0.3073] | +0.4583 [0.3705, 0.5450] |
| Deep intronic, sensitivity set | (a) without gene-level | 0.8277 [0.7596, 0.8974] | −0.0191 [−0.0426, 0.0012] | +0.2207 [0.1487, 0.2901] | +0.4754 [0.3795, 0.5711] |
| Non-canonical | (b) without DANN and GPN-MSA | 0.9197 [0.9021, 0.9359] | +0.0037 [−0.0012, 0.0088] | +0.1653 [0.1357, 0.1950] | +0.5559 [0.5149, 0.5934] |
| Donor region | (b) without DANN and GPN-MSA | 0.9692 [0.9552, 0.9816] | +0.0046 [−0.0001, 0.0104] | +0.1084 [0.0769, 0.1435] | +0.2814 [0.2315, 0.3329] |
| Acceptor region | (b) without DANN and GPN-MSA | 0.9504 [0.9181, 0.9763] | +0.0036 [−0.0051, 0.0132] | +0.0859 [0.0427, 0.1306] | +0.6018 [0.5157, 0.6804] |
| PPT region | (b) without DANN and GPN-MSA | 0.7746 [0.6793, 0.8641] | −0.0079 [−0.0358, 0.0182] | +0.0463 [−0.0413, 0.1320] | +0.6657 [0.5409, 0.7649] |
| Deep intronic | (b) without DANN and GPN-MSA | 0.8273 [0.7671, 0.8840] | +0.0032 [−0.0123, 0.0191] | +0.2763 [0.2076, 0.3441] | +0.4998 [0.4180, 0.5802] |
| Deep intronic, sensitivity set | (b) without DANN and GPN-MSA | 0.8484 [0.7875, 0.9121] | +0.0015 [−0.0140, 0.0162] | +0.2413 [0.1720, 0.3109] | +0.4960 [0.4018, 0.5913] |
| Non-canonical | (c) without both | 0.9065 [0.8865, 0.9252] | −0.0094 [−0.0169, −0.0029] | +0.1525 [0.1242, 0.1813] | +0.5427 [0.5033, 0.5795] |
| Donor region | (c) without both | 0.9600 [0.9407, 0.9763] | −0.0046 [−0.0130, 0.0032] | +0.0997 [0.0677, 0.1335] | +0.2722 [0.2258, 0.3201] |
| Acceptor region | (c) without both | 0.9491 [0.9165, 0.9741] | +0.0023 [−0.0121, 0.0168] | +0.0846 [0.0428, 0.1308] | +0.6005 [0.5173, 0.6737] |
| PPT region | (c) without both | 0.7134 [0.6051, 0.8263] | −0.0691 [−0.1211, −0.0234] | −0.0149 [−0.1080, 0.0867] | +0.6044 [0.4669, 0.7232] |
| Deep intronic | (c) without both | 0.8039 [0.7383, 0.8656] | −0.0201 [−0.0423, 0.0013] | +0.2529 [0.1873, 0.3182] | +0.4764 [0.3937, 0.5584] |
| Deep intronic, sensitivity set | (c) without both | 0.8347 [0.7659, 0.9005] | −0.0122 [−0.0337, 0.0057] | +0.2276 [0.1593, 0.2931] | +0.4823 [0.3906, 0.5761] |
| Non-canonical | (d) without both, gene-disjoint | 0.9030 [0.8834, 0.9214] | −0.0129 [−0.0215, −0.0045] | +0.1497 [0.1211, 0.1787] | +0.5393 [0.4986, 0.5765] |
| Donor region | (d) without both, gene-disjoint | 0.9582 [0.9399, 0.9736] | −0.0064 [−0.0162, 0.0033] | +0.0990 [0.0681, 0.1330] | +0.2704 [0.2219, 0.3189] |
| Acceptor region | (d) without both, gene-disjoint | 0.9440 [0.9103, 0.9707] | −0.0028 [−0.0177, 0.0103] | +0.0794 [0.0346, 0.1263] | +0.5954 [0.5111, 0.6705] |
| PPT region | (d) without both, gene-disjoint | 0.7441 [0.6382, 0.8524] | −0.0384 [−0.0922, 0.0085] | +0.0158 [−0.0769, 0.1207] | +0.6351 [0.5049, 0.7441] |
| Deep intronic | (d) without both, gene-disjoint | 0.7974 [0.7331, 0.8572] | −0.0267 [−0.0520, −0.0039] | +0.2464 [0.1819, 0.3122] | +0.4699 [0.3855, 0.5527] |
| Deep intronic, sensitivity set | (d) without both, gene-disjoint | 0.8265 [0.7590, 0.8936] | −0.0203 [−0.0445, −0.0005] | +0.2195 [0.1499, 0.2883] | +0.4742 [0.3781, 0.5680] |

Gene-level features: pLI, LOEUF and the ClinGen haploinsufficiency score. (d) is (c) retrained with genes held out, as in the gene-disjoint analysis. auPRC, area under the precision–recall curve; AUROC, area under the receiver operating characteristic curve; B/LB, benign or likely benign; CI, confidence interval; P/LP, pathogenic or likely pathogenic; PPT, polypyrimidine tract.

**Table S8. SpliceVarDB, within-region comparison**

Strict endpoint: splice-altering versus normal variants; broad endpoint: splice-altering or low-frequency splice-altering versus normal. auPRC and AUROC are given for MetaSplice / SpliceAI / CADD.

| Endpoint | Region | Variants (positives) | auPRC | AUROC | MetaSplice minus SpliceAI | MetaSplice minus CADD |
| --- | --- | --- | --- | --- | --- | --- |
| Strict | Canonical acceptor site | 3,132 (3,126) | 0.9996 / 0.9977 / 0.9971 | 0.8203 / 0.4522 / 0.3675 | +0.0019 [+0.0005, +0.0030] | +0.0025 [+0.0008, +0.0041] |
| Strict | Donor region | 1,700 (1,399) | 0.9904 / 0.9777 / 0.9571 | 0.9650 / 0.9344 / 0.8690 | +0.0128 [+0.0080, +0.0186] | +0.0332 [+0.0239, +0.0433] |
| Strict | Acceptor region | 1,121 (711) | 0.9667 / 0.9423 / 0.9109 | 0.9368 / 0.9175 / 0.8546 | +0.0248 [+0.0146, +0.0360] | +0.0559 [+0.0417, +0.0701] |
| Strict | PPT region | 1,246 (244) | 0.8128 / 0.8220 / 0.5484 | 0.9029 / 0.9276 / 0.7483 | −0.0092 [−0.0359, +0.0173] | +0.2644 [+0.2083, +0.3139] |
| Strict | Branch-point region | 1,175 (63) | 0.4492 / 0.3747 / 0.1588 | 0.7903 / 0.7852 / 0.6513 | +0.0747 [−0.0062, +0.1613] | +0.2904 [+0.1756, +0.3987] |
| Strict | Deep intronic | 2,157 (273) | 0.8700 / 0.8524 / 0.6204 | 0.9262 / 0.9269 / 0.8125 | +0.0172 [−0.0065, +0.0406] | +0.2497 [+0.1965, +0.3028] |
| Strict | Deep intronic, sensitivity set | 1,846 (163) | 0.8384 / 0.8173 / 0.5763 | 0.9093 / 0.9151 / 0.8071 | +0.0201 [−0.0112, +0.0506] | +0.2622 [+0.1886, +0.3309] |
| Broad | Canonical acceptor site | 3,883 (3,877) | 0.9997 / 0.9982 / 0.9976 | 0.8192 / 0.4466 / 0.3607 | +0.0015 [+0.0005, +0.0025] | +0.0020 [+0.0007, +0.0033] |
| Broad | Donor region | 2,675 (2,374) | 0.9820 / 0.9707 / 0.9613 | 0.8742 / 0.8503 / 0.7871 | +0.0114 [+0.0081, +0.0150] | +0.0206 [+0.0153, +0.0269] |
| Broad | Acceptor region | 2,151 (1,741) | 0.9347 / 0.9183 / 0.9122 | 0.7453 / 0.7428 / 0.6970 | +0.0170 [+0.0115, +0.0224] | +0.0225 [+0.0146, +0.0305] |
| Broad | PPT region | 2,891 (1,889) | 0.7430 / 0.7287 / 0.7072 | 0.5687 / 0.5732 / 0.5348 | +0.0147 [+0.0034, +0.0258] | +0.0358 [+0.0191, +0.0518] |
| Broad | Branch-point region | 2,894 (1,782) | 0.6215 / 0.6384 / 0.6422 | 0.4838 / 0.5234 / 0.5230 | −0.0167 [−0.0322, −0.0018] | −0.0207 [−0.0391, −0.0016] |
| Broad | Deep intronic | 5,038 (3,154) | 0.7071 / 0.6904 / 0.7091 | 0.5546 / 0.5578 / 0.5731 | +0.0164 [+0.0064, +0.0264] | −0.0020 [−0.0149, +0.0116] |
| Broad | Deep intronic, sensitivity set | 4,375 (2,692) | 0.6769 / 0.6653 / 0.6906 | 0.5357 / 0.5411 / 0.5640 | +0.0107 [−0.0009, +0.0211] | −0.0136 [−0.0284, +0.0022] |

The canonical donor site (one normal variant) is omitted. Pooled over regions (strict endpoint), auPRC was 0.9928 for MetaSplice, 0.9853 for SpliceAI and 0.9809 for CADD. At MetaSplice scores of 0.1, 0.2 and 0.5, sensitivity for splice-altering variants was 92.8%, 91.5% and 89.0% and the false-positive rate among normal variants 1.2%, 1.0% and 0.7%. auPRC, area under the precision–recall curve; AUROC, area under the receiver operating characteristic curve; B/LB, benign or likely benign; CI, confidence interval; P/LP, pathogenic or likely pathogenic; PPT, polypyrimidine tract.

**Table S9a. Likelihood ratios of the derived score intervals in the test set's calibration cohort**

Each score is assigned to its highest attained interval; exact 95% CIs.

| Review status | Tool | Interval | P/LP | B/LB | Likelihood ratio [95% CI] | Point estimate | CI |
| --- | --- | --- | --- | --- | --- | --- | --- |
| ≥ 1 star | MetaSplice | PP3 Very Strong | 209 | 2 | 6.74e+03 [1.45e+03, 9.1e+04] | meets | clears |
| ≥ 1 star | MetaSplice | PP3 Strong | 296 | 71 | 269 [188, 390] | meets | clears |
| ≥ 1 star | MetaSplice | PP3 Moderate | 33 | 120 | 17.7 [9.51, 31.8] | meets | clears |
| ≥ 1 star | MetaSplice | PP3 Supporting | 10 | 179 | 3.6 [1.31, 8.38] | meets | inconclusive |
| ≥ 1 star | MetaSplice | Indeterminate | 16 | 2,302 | 0.448 [0.226, 0.8] |  |  |
| ≥ 1 star | MetaSplice | BP4 Supporting | 2 | 2,041 | 0.0632 [0.00503, 0.268] | meets | clears |
| ≥ 1 star | MetaSplice | BP4 Moderate | 6 | 9,967 | 0.0388 [0.0119, 0.0931] | meets | clears |
| ≥ 1 star | MetaSplice | BP4 Strong | 6 | 22,581 | 0.0171 [0.00532, 0.0407] | meets | inconclusive |
| ≥ 1 star | SpliceAI | PP3 Strong | 418 | 129 | 209 [161, 271] | meets | clears |
| ≥ 1 star | SpliceAI | PP3 Moderate | 66 | 264 | 16.1 [10.6, 24] | meets | clears |
| ≥ 1 star | SpliceAI | PP3 Supporting | 36 | 455 | 5.1 [3.08, 8.08] | meets | clears |
| ≥ 1 star | SpliceAI | Indeterminate | 19 | 1,629 | 0.752 [0.4, 1.3] |  |  |
| ≥ 1 star | SpliceAI | BP4 Supporting | 5 | 2,612 | 0.123 [0.0321, 0.327] | meets | clears |
| ≥ 1 star | SpliceAI | BP4 Moderate | 32 | 31,872 | 0.0647 [0.042, 0.0947] | meets | clears |
| ≥ 1 star | SpliceAI | NoScore | 2 | 302 | 0.427 [0.0314, 1.97] |  |  |
| all statuses | MetaSplice | PP3 Very Strong | 217 | 2 | 6.8e+03 [1.47e+03, 9.15e+04] | meets | clears |
| all statuses | MetaSplice | PP3 Strong | 304 | 71 | 268 [187, 388] | meets | clears |
| all statuses | MetaSplice | PP3 Moderate | 33 | 120 | 17.2 [9.24, 30.9] | meets | clears |
| all statuses | MetaSplice | PP3 Supporting | 10 | 179 | 3.5 [1.27, 8.15] | meets | inconclusive |
| all statuses | MetaSplice | Indeterminate | 16 | 2,302 | 0.435 [0.22, 0.777] |  |  |
| all statuses | MetaSplice | BP4 Supporting | 2 | 2,041 | 0.0614 [0.00489, 0.26] | meets | clears |
| all statuses | MetaSplice | BP4 Moderate | 7 | 9,967 | 0.044 [0.0151, 0.0994] | meets | clears |
| all statuses | MetaSplice | BP4 Strong | 6 | 22,583 | 0.0166 [0.00517, 0.0395] | meets | inconclusive |
| all statuses | SpliceAI | PP3 Strong | 430 | 129 | 209 [161, 271] | meets | clears |
| all statuses | SpliceAI | PP3 Moderate | 68 | 264 | 16.1 [10.7, 23.9] | meets | clears |
| all statuses | SpliceAI | PP3 Supporting | 37 | 455 | 5.09 [3.1, 8.03] | meets | clears |
| all statuses | SpliceAI | Indeterminate | 19 | 1,629 | 0.73 [0.388, 1.26] |  |  |
| all statuses | SpliceAI | BP4 Supporting | 6 | 2,612 | 0.144 [0.0432, 0.353] | meets | clears |
| all statuses | SpliceAI | BP4 Moderate | 33 | 31,874 | 0.0648 [0.0424, 0.0943] | meets | clears |
| all statuses | SpliceAI | NoScore | 2 | 302 | 0.415 [0.0305, 1.91] |  |  |

The BP4 Strong level was not selected. The candidate rule combines the PP3 Strong and Very Strong intervals, and the BP4 Moderate and Strong intervals, at Supporting strength. 'Meets': the point estimate meets the requirement at a prior of 0.0441; 'clears': the CI bound meets it. auPRC, area under the precision–recall curve; AUROC, area under the receiver operating characteristic curve; B/LB, benign or likely benign; CI, confidence interval; P/LP, pathogenic or likely pathogenic; PPT, polypyrimidine tract.

**Table S9b. Codes under the candidate rule and the ClinGen SpliceAI rule, all configurations**

SpliceAI raw: SpliceAI 1.3.1 without masking over ±4,999 nucleotides, first record for the allele (primary) or largest value over all genes; SpliceAI masked: precomputed masked scores.

| Set | Rule | P/LP with PP3 | B/LB with PP3 | B/LB with BP4 | P/LP with BP4 | PP3 likelihood ratio | BP4 likelihood ratio |
| --- | --- | --- | --- | --- | --- | --- | --- |
| calibration cohort, ≥ 1 star | MetaSplice | 505 (87.37%) | 73 (0.20%) | 32,548 (87.35%) | 12 (2.08%) | 446 [331, 609] | 0.0238 [0.0111, 0.0441] |
| calibration cohort, ≥ 1 star | SpliceAI raw, first record | 524 (90.66%) | 1,654 (4.44%) | 32,966 (88.47%) | 39 (6.75%) | 20.4 [18.7, 22.2] | 0.0763 [0.0519, 0.107] |
| calibration cohort, ≥ 1 star | SpliceAI raw, all genes | 537 (92.91%) | 1,671 (4.48%) | 32,923 (88.35%) | 25 (4.33%) | 20.7 [19.1, 22.4] | 0.049 [0.0298, 0.0753] |
| calibration cohort, ≥ 1 star | SpliceAI masked | 520 (89.97%) | 848 (2.28%) | 34,999 (93.92%) | 38 (6.57%) | 39.5 [35.4, 44] | 0.07 [0.0474, 0.0989] |
| calibration cohort, all statuses | MetaSplice | 521 (87.56%) | 73 (0.20%) | 32,550 (87.35%) | 13 (2.18%) | 447 [332, 610] | 0.025 [0.0121, 0.0454] |
| calibration cohort, all statuses | SpliceAI raw, first record | 540 (90.76%) | 1,654 (4.44%) | 32,968 (88.47%) | 40 (6.72%) | 20.4 [18.7, 22.2] | 0.076 [0.0519, 0.107] |
| calibration cohort, all statuses | SpliceAI raw, all genes | 553 (92.94%) | 1,671 (4.48%) | 32,925 (88.35%) | 26 (4.37%) | 20.7 [19.1, 22.4] | 0.0495 [0.0304, 0.0754] |
| calibration cohort, all statuses | SpliceAI masked | 535 (89.92%) | 848 (2.28%) | 35,001 (93.92%) | 40 (6.72%) | 39.5 [35.4, 43.9] | 0.0716 [0.049, 0.1] |
| resolved VUS, ≥ 1 star | MetaSplice | 139 (90.85%) | 7 (0.99%) | 512 (72.52%) | 0 (0.00%) | 91.6 [38.3, 276] | 0 [0, 0.0412] |
| resolved VUS, ≥ 1 star | SpliceAI raw, first record | 145 (94.77%) | 220 (31.16%) | 455 (64.45%) | 8 (5.23%) | 3.04 [2.53, 3.59] | 0.0811 [0.0293, 0.179] |
| resolved VUS, ≥ 1 star | SpliceAI raw, all genes | 150 (98.04%) | 220 (31.16%) | 452 (64.02%) | 3 (1.96%) | 3.15 [2.66, 3.65] | 0.0306 [0.00458, 0.104] |
| resolved VUS, ≥ 1 star | SpliceAI masked | 144 (94.12%) | 113 (16.01%) | 559 (79.18%) | 7 (4.58%) | 5.88 [4.57, 7.49] | 0.0578 [0.0195, 0.131] |
| resolved VUS, all statuses | MetaSplice | 139 (90.85%) | 7 (0.99%) | 514 (72.60%) | 0 (0.00%) | 91.9 [38.4, 276] | 0 [0, 0.0411] |
| resolved VUS, all statuses | SpliceAI raw, first record | 145 (94.77%) | 220 (31.07%) | 457 (64.55%) | 8 (5.23%) | 3.05 [2.54, 3.6] | 0.081 [0.0292, 0.179] |
| resolved VUS, all statuses | SpliceAI raw, all genes | 150 (98.04%) | 220 (31.07%) | 454 (64.12%) | 3 (1.96%) | 3.16 [2.67, 3.66] | 0.0306 [0.00457, 0.104] |
| resolved VUS, all statuses | SpliceAI masked | 144 (94.12%) | 113 (15.96%) | 561 (79.24%) | 7 (4.58%) | 5.9 [4.58, 7.51] | 0.0577 [0.0195, 0.131] |

A variant without a score receives no code. auPRC, area under the precision–recall curve; AUROC, area under the receiver operating characteristic curve; B/LB, benign or likely benign; CI, confidence interval; P/LP, pathogenic or likely pathogenic; PPT, polypyrimidine tract.

**Table S9c. Differences in code shares between the candidate rule and the SpliceAI rule**

MetaSplice share minus SpliceAI share, in percentage points, with 95% CIs from class-stratified resampling and from gene resampling (1,000 resamples each).

| Set | Comparison | Share | Difference | Stratified CI | Gene-resampling CI |
| --- | --- | --- | --- | --- | --- |
| calibration cohort, ≥ 1 star | MetaSplice minus SpliceAI raw, first record | P/LP given PP3 | −3.29 | [−5.88, −0.87] | [−6.94, 0.98] |
| calibration cohort, ≥ 1 star | MetaSplice minus SpliceAI raw, first record | B/LB given PP3 | −4.24 | [−4.46, −4.03] | [−4.48, −4.00] |
| calibration cohort, ≥ 1 star | MetaSplice minus SpliceAI raw, first record | B/LB given BP4 | −1.12 | [−1.49, −0.74] | [−2.17, −0.16] |
| calibration cohort, ≥ 1 star | MetaSplice minus SpliceAI raw, first record | P/LP given BP4 | −4.67 | [−6.57, −2.77] | [−8.61, −1.54] |
| calibration cohort, ≥ 1 star | MetaSplice minus SpliceAI raw, all genes | P/LP given PP3 | −5.54 | [−7.79, −3.29] | [−7.74, −3.24] |
| calibration cohort, ≥ 1 star | MetaSplice minus SpliceAI raw, all genes | B/LB given PP3 | −4.29 | [−4.51, −4.08] | [−4.52, −4.04] |
| calibration cohort, ≥ 1 star | MetaSplice minus SpliceAI raw, all genes | B/LB given BP4 | −1.01 | [−1.39, −0.62] | [−2.04, −0.06] |
| calibration cohort, ≥ 1 star | MetaSplice minus SpliceAI raw, all genes | P/LP given BP4 | −2.25 | [−3.81, −0.87] | [−3.75, −0.72] |
| calibration cohort, ≥ 1 star | MetaSplice minus SpliceAI masked | P/LP given PP3 | −2.60 | [−5.19, 0.00] | [−5.43, 0.64] |
| calibration cohort, ≥ 1 star | MetaSplice minus SpliceAI masked | B/LB given PP3 | −2.08 | [−2.23, −1.94] | [−2.25, −1.92] |
| calibration cohort, ≥ 1 star | MetaSplice minus SpliceAI masked | B/LB given BP4 | −6.58 | [−6.93, −6.23] | [−7.48, −5.66] |
| calibration cohort, ≥ 1 star | MetaSplice minus SpliceAI masked | P/LP given BP4 | −4.50 | [−6.40, −2.77] | [−6.84, −2.48] |
| calibration cohort, all statuses | MetaSplice minus SpliceAI raw, first record | P/LP given PP3 | −3.19 | [−5.71, −0.67] | [−6.45, 1.15] |
| calibration cohort, all statuses | MetaSplice minus SpliceAI raw, first record | B/LB given PP3 | −4.24 | [−4.45, −4.03] | [−4.48, −4.00] |
| calibration cohort, all statuses | MetaSplice minus SpliceAI raw, first record | B/LB given BP4 | −1.12 | [−1.53, −0.74] | [−2.20, −0.14] |
| calibration cohort, all statuses | MetaSplice minus SpliceAI raw, first record | P/LP given BP4 | −4.54 | [−6.39, −2.86] | [−8.37, −1.60] |
| calibration cohort, all statuses | MetaSplice minus SpliceAI raw, all genes | P/LP given PP3 | −5.38 | [−7.73, −3.19] | [−7.68, −2.99] |
| calibration cohort, all statuses | MetaSplice minus SpliceAI raw, all genes | B/LB given PP3 | −4.29 | [−4.50, −4.07] | [−4.53, −4.03] |
| calibration cohort, all statuses | MetaSplice minus SpliceAI raw, all genes | B/LB given BP4 | −1.01 | [−1.42, −0.63] | [−2.03, −0.06] |
| calibration cohort, all statuses | MetaSplice minus SpliceAI raw, all genes | P/LP given BP4 | −2.18 | [−3.87, −0.84] | [−3.67, −0.87] |
| calibration cohort, all statuses | MetaSplice minus SpliceAI masked | P/LP given PP3 | −2.35 | [−4.87, 0.17] | [−5.24, 0.64] |
| calibration cohort, all statuses | MetaSplice minus SpliceAI masked | B/LB given PP3 | −2.08 | [−2.24, −1.93] | [−2.24, −1.92] |
| calibration cohort, all statuses | MetaSplice minus SpliceAI masked | B/LB given BP4 | −6.58 | [−6.93, −6.23] | [−7.53, −5.65] |
| calibration cohort, all statuses | MetaSplice minus SpliceAI masked | P/LP given BP4 | −4.54 | [−6.39, −2.86] | [−6.58, −2.56] |
| resolved VUS, ≥ 1 star | MetaSplice minus SpliceAI raw, first record | P/LP given PP3 | −3.92 | [−9.15, 0.65] | [−10.69, 3.19] |
| resolved VUS, ≥ 1 star | MetaSplice minus SpliceAI raw, first record | B/LB given PP3 | −30.17 | [−33.57, −26.77] | [−34.52, −26.37] |
| resolved VUS, ≥ 1 star | MetaSplice minus SpliceAI raw, first record | B/LB given BP4 | +8.07 | [4.25, 12.04] | [4.00, 12.37] |
| resolved VUS, ≥ 1 star | MetaSplice minus SpliceAI raw, first record | P/LP given BP4 | −5.23 | [−8.51, −1.96] | [−10.83, −0.80] |
| resolved VUS, ≥ 1 star | MetaSplice minus SpliceAI raw, all genes | P/LP given PP3 | −7.19 | [−11.76, −3.27] | [−12.50, −2.63] |
| resolved VUS, ≥ 1 star | MetaSplice minus SpliceAI raw, all genes | B/LB given PP3 | −30.17 | [−33.57, −26.77] | [−34.52, −26.37] |
| resolved VUS, ≥ 1 star | MetaSplice minus SpliceAI raw, all genes | B/LB given BP4 | +8.50 | [4.67, 12.33] | [4.37, 12.80] |
| resolved VUS, ≥ 1 star | MetaSplice minus SpliceAI raw, all genes | P/LP given BP4 | −1.96 | [−4.58, 0.00] | [−4.48, 0.00] |
| resolved VUS, ≥ 1 star | MetaSplice minus SpliceAI masked | P/LP given PP3 | −3.27 | [−8.50, 1.96] | [−10.12, 4.08] |
| resolved VUS, ≥ 1 star | MetaSplice minus SpliceAI masked | B/LB given PP3 | −15.01 | [−17.56, −12.32] | [−18.06, −12.07] |
| resolved VUS, ≥ 1 star | MetaSplice minus SpliceAI masked | B/LB given BP4 | −6.66 | [−10.06, −2.97] | [−10.25, −2.89] |
| resolved VUS, ≥ 1 star | MetaSplice minus SpliceAI masked | P/LP given BP4 | −4.58 | [−7.84, −1.31] | [−9.49, −0.79] |
| resolved VUS, all statuses | MetaSplice minus SpliceAI raw, first record | P/LP given PP3 | −3.92 | [−9.15, 0.65] | [−10.28, 3.60] |
| resolved VUS, all statuses | MetaSplice minus SpliceAI raw, first record | B/LB given PP3 | −30.08 | [−33.76, −26.69] | [−34.38, −26.31] |
| resolved VUS, all statuses | MetaSplice minus SpliceAI raw, first record | B/LB given BP4 | +8.05 | [4.38, 12.15] | [3.93, 12.44] |
| resolved VUS, all statuses | MetaSplice minus SpliceAI raw, first record | P/LP given BP4 | −5.23 | [−8.51, −1.96] | [−11.05, −1.20] |
| resolved VUS, all statuses | MetaSplice minus SpliceAI raw, all genes | P/LP given PP3 | −7.19 | [−11.76, −3.27] | [−12.07, −2.73] |
| resolved VUS, all statuses | MetaSplice minus SpliceAI raw, all genes | B/LB given PP3 | −30.08 | [−33.76, −26.69] | [−34.38, −26.31] |
| resolved VUS, all statuses | MetaSplice minus SpliceAI raw, all genes | B/LB given BP4 | +8.47 | [4.80, 12.57] | [4.41, 12.65] |
| resolved VUS, all statuses | MetaSplice minus SpliceAI raw, all genes | P/LP given BP4 | −1.96 | [−4.58, 0.00] | [−4.70, 0.00] |
| resolved VUS, all statuses | MetaSplice minus SpliceAI masked | P/LP given PP3 | −3.27 | [−8.50, 1.96] | [−10.06, 4.17] |
| resolved VUS, all statuses | MetaSplice minus SpliceAI masked | B/LB given PP3 | −14.97 | [−17.66, −11.86] | [−18.00, −12.04] |
| resolved VUS, all statuses | MetaSplice minus SpliceAI masked | B/LB given BP4 | −6.64 | [−10.31, −3.11] | [−10.29, −3.08] |
| resolved VUS, all statuses | MetaSplice minus SpliceAI masked | P/LP given BP4 | −4.58 | [−7.84, −1.31] | [−9.72, −0.75] |

auPRC, area under the precision–recall curve; AUROC, area under the receiver operating characteristic curve; B/LB, benign or likely benign; CI, confidence interval; P/LP, pathogenic or likely pathogenic; PPT, polypyrimidine tract.

**Table S10. Codes among the remaining variants of uncertain significance**

60,776 variants uncertain or conflicting on 15 April 2026, in the calibration cohort, and not classified P/LP or B/LB by 28 September 2026.

| Rule | PP3 | BP4 | Neither |
| --- | --- | --- | --- |
| MetaSplice | 14,939 (24.6%) | 21,482 (35.3%) | 24,355 (40.1%) |
| SpliceAI raw, first record | 23,508 (38.7%) | 31,179 (51.3%) | 6,089 (10.0%) |
| SpliceAI raw, all genes | 23,667 (38.9%) | 31,010 (51.0%) | 6,099 (10.0%) |
| SpliceAI masked | 19,785 (32.6%) | 36,705 (60.4%) | 4,286 (7.1%) |

MetaSplice scores of at least 0.1, 0.2 and 0.5: 33.1%, 30.0%, 25.1%. auPRC, area under the precision–recall curve; AUROC, area under the receiver operating characteristic curve; B/LB, benign or likely benign; CI, confidence interval; P/LP, pathogenic or likely pathogenic; PPT, polypyrimidine tract.
